# A new method based on genome alignments provides a highly resolutive target enrichment set for weevils (Coleoptera, Curculionoidea)

**DOI:** 10.64898/2026.05.09.724036

**Authors:** B. Zelvelder, L. Benoit, A. Loiseau, J. Haran, R. Allio

**Author notes:** Correspondence to be sent to: Benjamin Zelvelder, ISEM, Univ Montpellier, CNRS, IRD, Montpellier, France; Rémi Allio, Centre de Biologie pour la Gestion des Populations, CIRAD, INRAE, IRD, Institut Agro, Univ. Montpellier, Montpellier, France.

## Abstract

Target enrichment methods have provided unprecedented advances in phylogenomics. Targeting hundreds of conserved regions has proven to be a good tradeoff between cost and efficiency, while being useful for museomics and diversified non-model clades. Unfortunately, current methods used for identifying such regions involve high degrees of conservation within targeted elements, usually pushing researchers to rely on flanking data with little guarantee for homology. With a growing number of high quality genomes available throughout the Tree of Life emerges new opportunities to improve marker selection. In this study, we introduce GABBI, a new method for designing target capture probes by taking advantage of genome alignments, avoiding the selection of a single reference genome that can cause notable biases. We compare GABBI-derived markers to the most commonly used probe design method, PHYLUCE, at two taxonomic scales, the weevil superfamily Curculionoidea and the tribe Pachyrhynchini. At both taxonomic scales, results show that our new method allows identifying more variable loci that prove to be more phylogenetically resolutive than the PHYLUCE-derived ones. Doing so, we provide the first probe set specifically designed for weevils, targeting a wide set of 4,255 shared homologous regions, encouraging future research on systematics and macroevolution of one of the most diverse and economically important groups of insects. By providing GABBI as an automated and open-access pipeline, we hope to open new probe design opportunities to other taxonomic groups that face similar phylogenetic obstacles.

## TARGETED SEQUENCING IN THE INTEGRATIVE TAXONOMY ERA

Reconstructing the Tree of Life was once a dream that is rapidly becoming real thanks to the never ending progress of computational power and sequencing technologies. However, despite the exponential growth of molecular datasets, many phylogenomic studies continue to face topological uncertainty, not only because of hard to resolve biological events (e.g. ILS, HGT, hybridization etc.) but also because of methodological limits linked to phylogenetic models or genetic markers (see the reviews of Kapli et al. 2020; Fleming et al. 2023; Steenwyk et al. 2023). Although analyses limited to only one or a few genetic markers are being continuously disputed by larger molecular datasets, an increasing number of genetic markers doesn’t necessarily allow for better species tree reconstruction (Delsuc et al. 2005; Jeffroy et al. 2006; Philippe et al. 2011; Salichos and Rokas 2013; Shen et al. 2017). Instead, marker selection plays a key role in phylogenomic reconstructions, as different markers vary considerably in their informativeness, evolutionary rate, and susceptibility to systematic biases (Townsend 2007; López-Giráldez et al. 2013; Mongiardino Koch 2021). However, marker selection often requires access to multiple reference genomes. Although sequencing high quality genomes has become increasingly accessible, notably owing to flourishing international consortium (ERGA, i5k, DTOL), it is still far from being routinely used due to its high cost and difficult application to non-model species and very diverse taxa. Thankfully, harvesting genome wide markers can be achieved with genome reduction methods like targeted sequencing. Target capture methods help tackle technical issues linked to whole genome sequences (WGS) by providing large scale genomic markers at low cost while being suitable for large datasets of non model species and museum specimens (McCormack et al. 2013; Van Dam et al. 2017; Derkarabetian et al. 2019; Houston et al. 2022). Multiple methods of targeted enrichment have emerged during the last two decades such as ultraconserved elements (UCE; Faircloth et al. 2012), anchored hybrid enrichment (AHE; Lemmon et al. 2012), HyRAD-X (Toussaint et al. 2021), or whole genome enrichment (WGE; Carpenter et al. 2013). All of these methods have in common that they rely on conserved elements – often chosen to be anchored in single copy genomic regions with relevant phylogenetic signal – to capture targeted sequences with hybridization probes used during target enrichment protocols. The main differences being how targeted regions are identified and how many genetic markers can be targeted by the final probe set.

Thanks to the emergence of these different methods, most major taxonomic groups have one or multiple probe sets available for target capture, like amniotes (Faircloth et al. 2012), vertebrates (Lemmon et al. 2012), angiosperms (Johnson et al. 2019), and many orders of arthropods (Faircloth 2017; Zhang et al. 2023; Jasso-Martínez et al. 2025; Cannizzaro and Berg 2026). Nevertheless, a growing part of literature shows that probe sets designed at wide taxonomic scales don’t meet the high degree of homology and variability needed to solve recalcitrant phylogenetic nodes (Van Dam et al. 2017; Lee et al. 2021; Baca et al. 2023; Haran et al. 2023b). Selected markers can either be too conserved, pushing the phylogenetic signal to flanking regions that can be difficult to align, because farther from the conserved element, sequences rapidly become too variable and poorly homologous. Coding markers are often saturated at large taxonomic ranges, and designing probes on very few representative taxa can bias marker selection and reduce marker recovery in lineages that were not represented during probe design. To bypass these limits, custom probe sets are increasingly used at shallower taxonomic scales to identify more numerous and variable markers, although restricting their use to one or a handful of studies (e.g. Branstetter et al. 2017; Yardeni et al. 2022; Van Dam et al. 2023; Gutiérrez-Trejo et al. 2024).

## DESIGNING PROBES IN THE GENOMIC ERA

Most of the limits that have been identified in target capture approaches can be associated with marker selection. Few biases persist in the techniques commonly used to identify conserved elements and to design the probes to target them. Firstly, probe sets designed at a broad taxonomic scale rely on a very limited number of reference taxa selected to represent tens of thousands of species that diverged millions of years ago (e.g. Faircloth et al. 2012, 2015; Lemmon et al. 2012). For this reason, genetic markers have to be highly conserved to be found confidently at those evolutionary scales, leading to, on one hand, targeted markers that include few informative sites and, on the other hand, a great difficulty in aligning flanking regions of very divergent taxa. Incidentally, methods used to find these conserved elements have to rely on mapping genetic elements on a given genome, which also implies a high level of conservation and reinforces the selection of markers that are uninformative and saturated without flanking data. Lastly, those methods all rely, at some point and to our knowledge, on a single genomic reference, restricting the number of potential loci that can be identified in the first place.

In this study, we introduce a novel approach to design target capture probe sets that attempts to tackle these biases. We designed a pipeline based on a genome alignment that offers the advantage of being free from the ‘single reference’ bias, allowing the identification of shared, single copy homologous regions (hereafter referred to as SHR) that are no longer necessarily highly conserved. This approach takes advantage of the growing number of available genomes throughout the Tree of Life, thus providing a better representation of taxon variability during the probe design process. The entire pipeline for the Genome Alignment Bait Inference pipeline (GABBI) is openly available on github (https://github.com/bjzelvelder/GABBI.git).

## DESIGNING THE WEEVIL PROBE SET

Here, we follow this new GABBI approach of designing probe sets for the beetle superfamily Curculionoidea, also known as weevils. Weevils are one of the most economically important groups of insects, mainly known for their antagonistic relationship with plants (Zimmerman 1991; Anderson 1995), but also for their widely underrated pollination service in tropical and subtropical areas (Haran et al. 2023a). Represented by more than 62,000 described species worldwide and probably more than 200,000 awaiting description (Oberprieler et al. 2007), the weevils’ phylogeny is far from being completely resolved due to their rapid radiation during the late cretaceous period (McKenna et al. 2009). Great advances have been made recently thanks to targeted enrichment, notably the AHE probe set made available by Haddad et al. (2018), however, the ultradiverse and rapidly evolving weevils have raised shortcomings in phylogenomic analyses that clearly underlined the need for a specific weevil probe set capable of resolving the rapidly evolving nodes (Shin et al. 2018; Haran et al. 2023b; Li et al. 2023; Zelvelder et al. 2026).

In this study, we follow the GABBI approach to develop a new set of target enrichment probes specific to the weevil superfamily Curculionoidea. We compare this method with the most commonly used method for probe design, following the PHYLUCE pipeline, with *in silico* analyses at two taxonomic scales: the weevil superfamily (Coleoptera, Curculionoidea) and the easter-egg weevils tribe (Pachyrhynchini). We synthesized our new weevil probe set and tested it on a small yet representative batch of 12 weevil specimens conserved in dry collections or pure ethanol to evaluate its efficiency both *in silico* and *in vitro*.

## MATERIAL AND METHODS

### Study Overview

Two methods of probe design are being tested. The first and most commonly used method, based on the mapping of simulated reads on a chosen reference, will be referred to as “PHYLUCE” as it follows the pipeline of the same name (Faircloth 2016). The second method, newly established in this study, is a Genome Alignment Based method of Bait Inference referred to as its acronym GABBI. Both methods are tested on the same genomic datasets consisting of 56 whole genome sequences available on NCBI and 4 genomes that were newly generated in this study (Table S1; S2). Both methods follow a similar two step process. The first step consists in the identification of shared loci to form a temporary probe set. It is conducted on 9 chromosome-level genomes of Curculionidae available at the time of this study (Fig. 1a). The second step consists in the *in silico* validation of temporary probes on a larger genomic dataset to form the final probe set. It is conducted on 42 Curculionoidea genomes that represent most of the diversity of the Curculionoidea superfamily, including the nine initial genomes (Fig. 1b). Genomes that were newly generated in this study are included in this set of 42 genomes, three of which as assemblies (Table S1) and one as unassembled raw read data. Final probe sets are then used for *in silico* phylogenomic analyses at two taxonomic scales: the Curculionoidea scale, represented by 60 genomes including the ones used during probe design, hereafter named the *Curculionoidea dataset* (Table S2), and the tribe scale using 82 raw Pachyrhynchini target capture data from Van Dam et al. (2023), hereafter named the *Pachyrhynchini dataset*.

**Figure 1:**
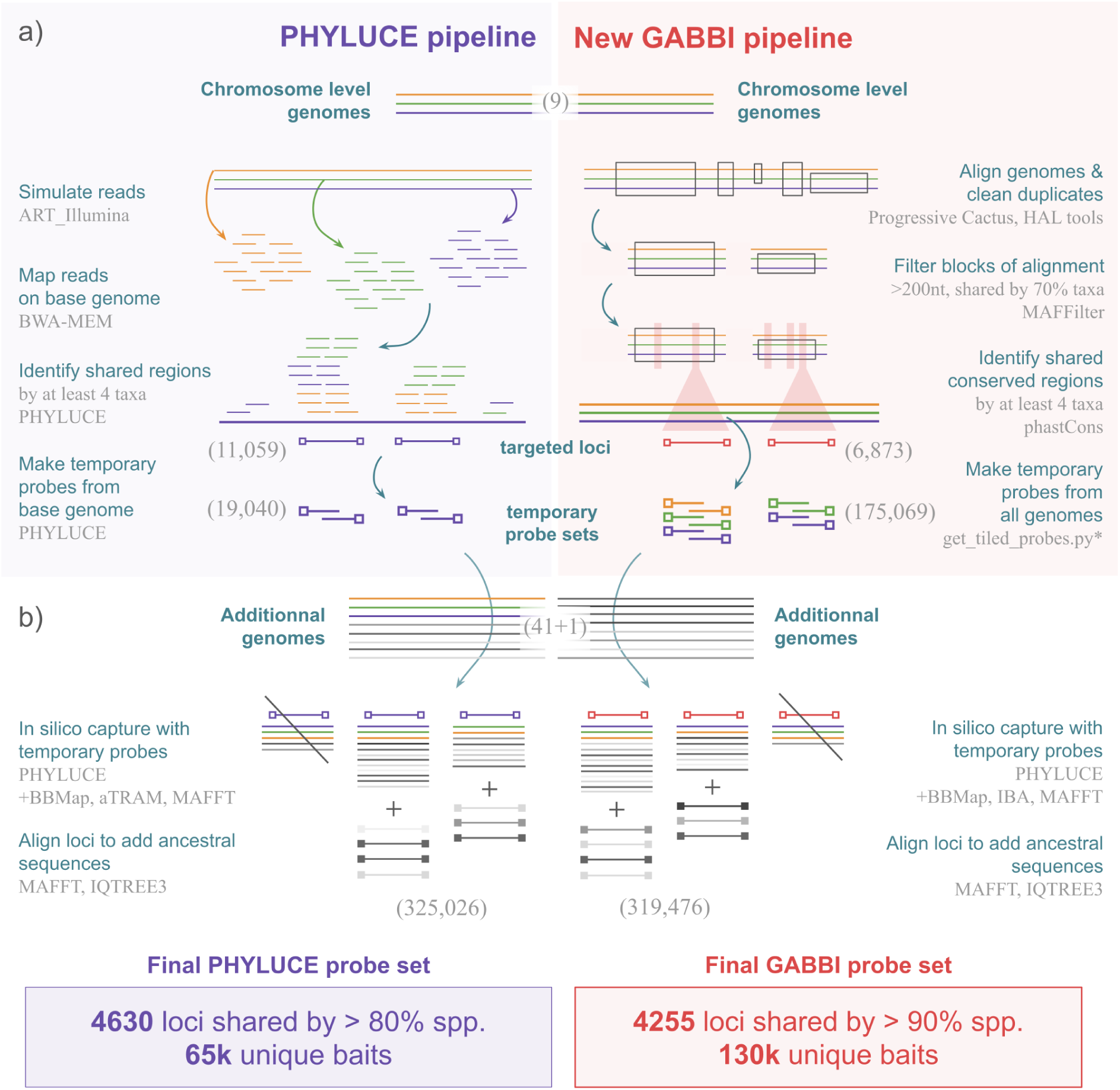
Step by step comparative flowchart of PHYLUCE and GABBI pipelines. PHYLUCE = Name of Faircloth’s pipeline (2016), GABBI = Genome Alignment Based Bait Inference, developed in this study. a) The first step, where both pipelines differ the most, consists in the identification of targeted loci from a set of chromosome-level genomes to make temporary probes. b) The second step, identical in both pipelines, consists in the validation of the temporary probe set on a set of additional genomes to select the final set of targeted loci. Resulting loci were handed over to myBaits^®^ for actual probe design and filtration. Numerical results and method names written in grey are specific to this study.

### Probe Design based on Read Mapping

Unless otherwise stated, designing the PHYLUCE probe set followed the aforenamed pipeline and scripts made available by Faircloth (2016; https://phyluce.readthedocs.io/en/latest/tutorials/tutorial-4.html). Reads of chromosome-level genomes were simulated with ART illumina (Huang et al. 2012) then mapped to the genome of *Polydrusus cervinus* with BWA-MEM (Li 2013). This base genome was chosen following Gustafson et al. (2019) recommendations as it minimizes the genetic distance with other considered genomes. From the coverage of mapped reads onto the base genome, conserved regions were identified and used to build a temporary probe set. These probes were then used to make *in silico* capture on the 41 assembled genomes (see the *Study Overview* section) and design a second set of probes. The latter was used to conduct the *in silico* capture of raw read data from the unassembled genome of *Isorhynchus* (see the *In Silico Target Capture* section).

The assembled target capture data was then shortened to the region targeted by other probes and resulting sequences added to the PHYLUCE pipeline with custom scripts (see *pipeline_phyluce.sh* on Zenodo at xxx). In order to match approximately with the number of loci obtained with GABBI, loci shared by at least 80% of taxa make up the final set of PHYLUCE targeted loci. In an attempt to better map the inner variation at the targeted loci and improve *in silico* or *in vitro* capture of taxa missing from our dataset, targeted loci were aligned and cleaned with *Phylomera* (see the *Phylogenomic analyses* section) to generate gene trees of each locus and infer ancestral sequences using the IQTREE3 option *-asr* (Wong et al. 2025). All targeted loci (before alignments) and their ancestral sequences were sent to myBaits^®^ for probe design, quality control and reduction by clustering redundant probes.

### Probe Design based on Genome Alignments

To design the GABBI probe set, we started with the genome alignment of the 9 chromosome-level genomes using Progressive Cactus (Armstrong et al. 2020). Resulting blocks of alignments were treated with HAL tools (Hickey et al. 2013) to filter out non-orthologous and highly repeated regions and filtered with MafFilter (Dutheil et al. 2014) to only keep blocks of alignments longer than 200 nucleotides and shared by more than 70% of extant and ancestral genomes inferred by cactus during the alignment process. Conserved regions within those blocks of alignment were identified using phastcons (Hubisz et al. 2011) on each extant genome and cross-blasted together with BLASTn (Altschul 1997). Regions found in single or low copy (in at most one lineage) and shared by at least four taxa were kept for the temporary set of loci. Ancestral sequences were then generated to enrich this set of targeted loci (see the *Probe Design based on Read Mapping* section) before making temporary probes with a custom script that mimics the behavior of the corresponding PHYLUCE command (see *pipeline_cactus.sh* on Zenodo). *In silico* capture of the 41 assembled genomes of weevils was conducted with PHYLUCE and completed with the target capture data from the *Isorhynchus* genome using IBA (Breinholt et al. 2018) instead of aTRAM (see the *In Silico Target Capture* section). At last, loci shared by at least 90% of taxa make up the final set of GABBI targeted loci. As with the PHYLUCE probe design, ancestral sequences were generated for each targeted locus before giving targeted loci to myBaits^®^ for final probe design and quality checks. The GABBI pipeline was further improved and automated in an easily usable singularity image openly available on github (https://github.com/bjzelvelder/GABBI.git).

### In Silico Target Capture

To compare PHYLUCE and GABBI probe design methods we simulated *in silico* target capture at two different taxonomic scales using the *Curculionoidea dataset* and the *Pachyrhynchini dataset* (see the *Study Overview* section). Unless otherwise stated, data processing was conducted using custom scripts written in *bash*, *python* or R (all of which are available on Zenodo).

#### Curculionoidea dataset

Reads of already assembled genomes were simulated using ART illumina with default error rate parameters and cleaned with fastp (Chen et al. 2018) while unassembled reads of *Isorhynchus* were used as is. To reduce the number of reads to process, BBMap (Bushnell 2014) was used to subsample reads based on a soft threshold (50%) of alignment identity with probe sequences, roughly mimicking hybrid enrichment. The assemblage of targeted markers was then conducted iteratively based on all probe sequences of each locus with a slightly modified version of aTRAM (Allen et al. 2015; singularity image on Zenodo), that runs SPAdes 4.2.0 (Prjibelski et al. 2020) with the following options to retrieve better results on low coverage target enrichment data: *--evalue 1e-3 --word-size 11 --blast-max-target-seqs 100 --spades-careful*. All sequences assembled from probes targeting the same locus are aligned with MAFFT v7.453 (Katoh 2005; Katoh and Standley 2013) to make a majority rule consensus of assembled sequences. To eliminate potential contaminations, gene duplication, and assembly errors, sequences that differ by more than 5% are filtered out if they make up less than 10% of sequences. Otherwise, the alignment is discarded and the marker is flagged. Markers that were flagged in more than two taxa are blacklisted and removed from the dataset as they may be duplicated in more than two lineages. Consensus sequences of all taxa are grouped under one multifasta file per marker for subsequent phylogenomic analyses.

#### Pachyrhynchini dataset

We downloaded the raw read data from the targeted sequencing of Van Dam et al. (2023) dataset that used a very large probe set targeting more than 10k loci, in an attempt to capture overlapping markers between our probe sets and theirs or with potential off-target reads. We followed the same process as the *Curculionoidea dataset in silico* target capture using fastp, BBMap, aTRAM and custom scripts for postprocessing. We also conducted the *in silico* target capture and phylogenomic analysis of this dataset using Van Dam et al.’s probe set to make our results comparable to theirs.

At both taxonomic scales, we also simulated *in silico* target capture using two currently available probe sets compatible with weevils such as the Coleoptera UCE set from Faircloth et al. (2017) and the Coleoptera AHE set from Haddad et al. (2018) following the same process described above.

### Phylogenomic Analyses

In this study, each dataset is analysed in a maximum likelihood (ML) framework carefully partitioned to improve phylogenomic inferences. To do so, we first need to recover reference sequences of each locus to separate core and flanking regions.

Reference sequences were generated for each marker by making a IUPAC consensus of the targeted regions alignments, filtered with HMMCleaner (Di Franco et al. 2019) and trimmed with AMAS (Borowiec 2016) to remove columns with more than 95% gaps. Resulting reference sequences are then combined into a single multifasta file easier to handle. The PHYLUCE and GABBI targeted loci were also annotated as ‘coding’ or ‘non-coding’ in a so-called annotation file to make a codon-wise partitioning of coding sequences (CDS). This information can be recovered using the coordinates of each targeted region in each genome. If a given marker overlaps with a CDS of at least one of the six annotated genomes of the dataset, it is flagged as ‘coding’ and treated accordingly as detailed below (see *pipeline_annotation.sh* on Zenodo for more details on annotating markers).

In order to make our phylogenomic analyses of target capture data more accessible, easier to reproduce and less time consuming, we developed *Phylomera*, a fast and versatile pipeline working in parallel (with GNU parallel; Tange 2011) to process any number of genomic markers from unaligned multifastas to phylogenomic tree inferences. Reference sequences and annotation files are given to *Phylomera* 0.8.3 with unaligned sequences of each marker to be processed. In this pipeline, sequences of each marker are first aligned with the MAFFT L-INS-I model. Reference sequences are split with SeqKit 2.9.0 (Shen et al. 2016) and aligned to each corresponding marker alignment using MAFFT option *--addfragments* to keep the integrity of the reference in the alignment (Katoh and Frith 2012). Based on reference coordinates in each alignment, alignments are split into core and flanking regions with AMAS *trim* option. The core region of all markers listed as CDS in the annotation file are processed with the OMM_MACSE v11.05 pipeline, as it is optimised for aligning and cleaning coding sequences and keeps a consistent open reading frame throughout coding sequences (Ranwez et al. 2018; Scornavacca et al. 2019). Otherwise, flanking regions, core regions annotated as non-coding and coding regions that retrieved no result after the OMM_MACSE pipeline are processed with HMMCleaner to filter non homologous regions and trimmed with AMAS to remove columns with more than 50% gaps. Sequences with more than 50% gaps are then removed from all alignments with *drop_short_seq.py* to limit biases linked to missing data. Alignments with more than 50%, 70% and 90% taxa represented in each dataset are concatenated into three supermatrices along with partition files composed of one partition per flanking region, core region and codons if coding markers are provided. ML species trees are then conducted for each supermatrix with IQTREE3 using the options *-m MFP+MERGE -rclusterf 10* that uses ModelFinder and PartitionFinder2 to estimate evolution models and merge partitions (Kalyaanamoorthy et al. 2017; Lanfear et al. 2017). As recommended by IQTREE authors for phylogenomic datasets, we included a perturbation level of heuristics of 0.2 using the *-pers* option. Bootstraps are estimated with ultrafast bootstraps (UFBS; Hoang et al. 2018) with *-bnni* option and Shimodaira-Hasegawa approximate likelihood-ratio test (SH-aLRT; Guindon et al. 2010). Gene tree inferences were also conducted with IQTREE3 for each marker without partitions to correctly estimate the alpha parameter of the Gamma distribution using a general time reversible model with eight Gamma parameters (GTR+G8) and allow a fair comparison between their median values (which is less biased by outliers than the mean). Coleoptera UCE and AHE probe sets couldn’t be annotated as coding or non-coding sequences. Comparative analyses between GABBI and these probe sets were therefore performed following the same process but did not consider coding and non-coding regions.

### In vitro Target Capture of Curculionoidea Specimen

The newly generated GABBI weevil probe set was tested *in vitro* on 12 specimens sampled between 2017 and 2025 and conserved in alcohol or in dry collections at CIRAD (Montpellier, France). DNA was extracted non-destructively following the instructions from the EZ-10 Spin Column Animal Genomic DNA Miniprep Kit (BioBasic) with an overnight lysis. Extracted DNA was then mechanically fragmented to 300-600 bp fragments using a Bioruptor Pico (Diagenode; 15s ON, 90s OFF, eight cycles). Libraries were built using a slightly modified protocol from the NEBNext Ultra II DNA Library Prep Kit for Illumina (New England BioLabs): all reagents volumes have been reduced by half, sizing was performed using a 0.7X ratio of AMPure XP beads (Beckman), indexing was done using 12 Unique Dual Index Primer Pairs of NEBNext Multiplex Oligos for Illumina (New England BioLabs) with 6 to 15 PCR cycles depending on the DNA yield. Libraries were then pooled together on an equimolar ratio to be enriched with GABBI weevil probe set, according to myBaits Custom Hybridization Capture Kit (Daicel Arbor Biosciences), with a probe amount reduced to 3.5 µL instead of 5.5 µL, and 2 final enrichment reactions of 12 PCR cycles. 52Go of libraries were sequenced by Novogene on a 10B lane (NovaSeq X, Illumina). Raw read data were cleaned with TRIMMOMATIC (Bolger et al. 2014). Trimmed reads then followed the same pipeline as other datasets, from SHR extraction using aTRAM and custom scripts to phylogenomic analyses with *Phylomera*.

### Comparative Statistics acquisition

Descriptive statistics of probe design and target capture results are estimated using command-line tools (e.g. *grep*, *awk*, *wc* etc.) based on resulting files or summary files generated with AMAS (see *stats_final.sh* on Zenodo). Markers annotated as CDS that passed the OMM_MACSE pipeline were split into codons to compute codon-wise statistics using AMAS *split* and *summary* options. Phylogenomic analyses statistics, such as tree lengths, bootstrap support and supermatrix size are taken from IQTREE output files. Other statistics, such as the proportion of parsimony informative sites (PPIS), the number of markers per taxa and size of core and flanking regions are harvested using AMAS *summary* option. PPIS distributions are compared with Wilcoxon-Mann-Whitney tests corrected for multiple comparisons with the Benjamini-Hochberg false-rate discovery in R v4.3.1 using the *rstatix* package (Kassambara 2023; R Core Team 2023).

## RESULTS

### Probe design

Both probe design methods provided thousands of candidate loci following the first mapping or genome-alignment step and had to be filtered. To avoid being too conservative at this early stage of probe design, genomic regions shared by at least 4 taxa out of 9 were kept for subsequent analyses. This resulted in 11,059 loci mapped on the base genome of *Polydrusus cervinus* using the PHYLUCE pipeline and 6,873 blocks of aligned loci with the GABBI pipeline. Translated into 2X tiled probe sequences, the PHYLUCE temporary probe set consists of 19,040 probes whereas the GABBI temporary probe set is made of 175,069 probes, which is much bigger because it is based on all 9 chromosome-level genomes instead of one. The temporary probe set is then tested on a bigger sample of 42 genomes to retain loci that are still represented in the largest share of taxa while maximizing the number of final loci.

Although unassembled, the lesser quality genome of *Isorhynchus* was interesting to add to the probe design to provide more variation in the Ceutorhynchinae clade that is hard to locate among true weevils, showing in the meantime that probe design can be enriched with genomes of lesser quality. The validation of the GABBI temporary probe set allowed us to keep 4,255 targeted loci shared by 90% taxa. Based on this result, we aimed to keep a similar number of loci from the PHYLUCE temporary probe set and could obtain 4,631 targeted loci shared by 80% taxa, deliberately surpassing the amount of GABBI loci to test its robustness given this small disadvantage (choosing 90% would have dropped the number of PHYLUCE targeted loci to only 2600). Based on these 42 genomic samples and ancestral sequences simulated at each locus, 325,026 sequences of targeted loci were obtained for PHYLUCE and 319,476 sequences for GABBI. From these targeted loci, the final probe sets were generated and filtered to remove redundant probes with over 75% overlap and 80% identity, which resulted in 4,630 loci targeted by 65,713 probes for the PHYLUCE probe set (median of 13 probes per marker) and 4,255 loci represented by 130,127 probes for the GABBI probe set (median of 22 probes per marker). The annotation of these loci estimated that 2,644 out of 4,630 PHYLUCE loci could be annotated as CDS, whereas 4,157 out of 4,255 GABBI loci could be annotated as CDS. Targeted loci of the PHYLUCE probe set are on average 211 bp long, ranging from 172 to 627 bp, while the targeted loci of the GABBI probe set are on average 237 bp long, ranging from 176 to 1249.

### In silico *target capture*

#### Curculionoidea dataset

In both probe sets, *in silico* target capture at the super-family scale provided high levels of marker recovery with a median number of 3,955 UCE (85.4%) for PHYLUCE and 3,727 SHR (87.6%) for GABBI (detailed in Table S2). In total, all markers were recovered in both probe sets, but blacklisting potentially duplicated markers slightly reduced the total number of markers to 4,466 for PHYLUCE and 4,136 for GABBI. Before alignments, assemblies were on average 1201.97 nt long for PHYLUCE and 1351.71 nt long for GABBI. Once each marker was aligned and thoroughly cleaned from non-homologous regions and columns with more than 50% ambiguous characters, core and flanking regions dropped to an average of 178.68 nt long and 179.21 nt long for PHYLUCE, and an average of 182.78 nt long and 277.32 nt long for GABBI, respectively. Average proportions of variable sites reach 0.646 in the PHYLUCE supermatrix and 0.640 in the GABBI supermatrix, while average proportions of parsimony informative sites reach 0.510 and 0.525, respectively. GABBI-derived alignments show significatively higher PPIS in core regions but not in flanking regions at this taxonomic scale (Fig. 2). As expected for coding regions due to the redundancy of the genetic code, PPIS at the core regions annotated as CDS are lower for the first and second codon positions rather than the third. Flanking regions (that are not annotated) also show higher PPIS compared to unannotated (probably non-coding) core regions, but this may only reflect the presence of coding regions in flanking data. Every pair-wise comparison between datasets has a *p-value* < 0.001.

**Figure 2:**
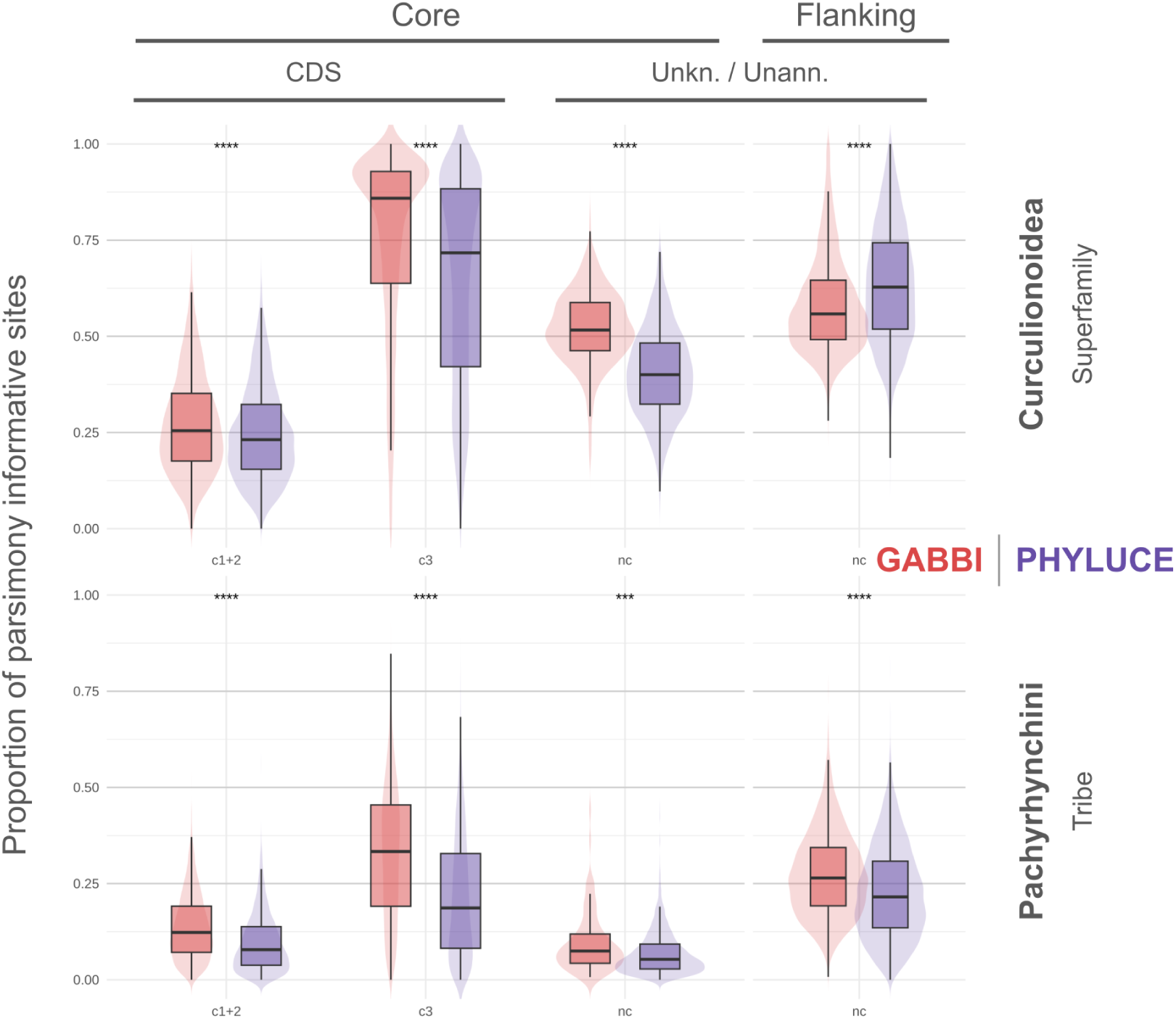
Proportions of parsimony informative sites between PHYLUCE and GABBI datasets. Markers were splitted into core and flanking regions and analysed accordingly. Core regions annotated as coding were splitted into codon positions 1+2 (pos 1+2) and 3 (pos 3) separately, while non-coding core regions and unannotated flanking regions were left untouched. For each alignment, a proportion of parsimony informative sites (PPIS) is calculated, their distribution in each subcategory is represented with a boxplot and a background violin plot. Distributions of PPIS values of GABBI-derived alignments are shown on the left and PHYLUCE-derived alignments on the right of each pair of boxplots. Asterisks indicate significance levels of pairwise mean comparisons using Wilcoxon-Mann-Whitney tests corrected for multiple comparisons using Benjamini-Hochberg false-rate discovery: *** p < 0.001, **** p < 0.0001.

#### Pachyrhynchini dataset

Even though we could not produce a sufficiently large dataset at the tribe scale with genomes available at the time of this study, we took the opportunity to try *in silico* target capture on the Pachyrhynchini dataset from Van Dam et al. (2023). Eventually, we were able to recover a median number of 3,122 markers per taxa with the PHYLUCE probe set, ranging from 103 to 4,529 per individual, and a median number of 2,946.5 markers per taxa with the GABBI probe set, ranging from 104 to 4,151 per individual. Blacklisted markers are more numerous in both probe sets (345 for PHYLUCE and 496 for GABBI) compared to the *Curculionoidea* dataset as it has potentially more contaminations and overall lesser quality data from true capture samples. Before alignments, assemblies were on average 394.965 bp long for PHYLUCE and 402.267 bp long for GABBI. Following alignment cleaning, core and flanking regions were on average 195.723 nt long and 79.098 nt long for PHYLUCE, and on average 210.555 nt long and 75.242 nt long for GABBI, respectively. Average proportions of variable sites reach 0.304 in the PHYLUCE supermatrix and 0.380 in the GABBI supermatrix, while global proportions of parsimony informative sites reach 0.157 and 0.227, respectively. GABBI-derived alignments show significatively higher PPIS in both core and flanking regions (Fig. 2). Unannotated core regions (probably non-coding) show very reduced PPIS in this dataset compared to the superfamily scale, but unannotated flanking regions still have intermediate PPIS that average around 0.227 for PHYLUCE and 0.274 for GABBI. PPIS distributions between the third codon position and the first two, as well as between unannotated core and flanking regions agree with the *Curculionoidea* dataset and are also significatively different at the *Pachyrhynchini* taxonomic scale (*p* < 0.0001; Fig. 2). In comparison, following the same methodological framework but using Van Dam et al.’s probe set of 12,406 custom UCE, we could use 11,852 UCE cleaned alignments with an average marker size of 380.656 bp, average PVS of 0.444 and average PPIS of 0.252.

### Phylogenomic tree inferences

#### Curculionoidea dataset

Alignments obtained from *in silico* capture at the superfamily scale were concatenated into supermatrices with three levels of missing data, consisting of markers with at least 50%, 70% or 90% of taxa represented. Supermatrix sizes range from 87,264 nt to 1,978,183 nt in the PHYLUCE dataset, and from 305,900 nt to 2,505,840 nt in the GABBI dataset. Overall, GABBI-derived trees performed better than PHYLUCE-derived ones, exhibiting less unsupported nodes and longer branch lengths (Fig. 3b; Table S3). Phylogenomic trees obtained show overall similar topologies that always agree on the position of *‘early diverging lineages’* (sensu Haran et al. 2023b), and the monophyly of both CEGH and CCCMS clades (Cyclominae, Entiminae, Gonipterini, Hyperinae, and Curculioninae, Conoderinae, Cossoninae, Molytinae, Scolytinae, respectively). Topologies mainly differ on the position of the ‘Curculioninae II’ clade within CCCMS (regrouping *Orchestes* and *Anthonomus genera;* Fig. 3a-b; S1-2). The total tree length of the PHYLUCE-derived and GABBI-derived trees represented in figure 3 (70% completeness) are respectively 8.771 and 12.834 units long and have respectively 5 and 1 unsupported nodes. Gene trees computed on all markers regardless of their completeness using GTR+G8 models allowed us to get a median value for the gamma function’s alpha parameter of 0.379 for PHYLUCE markers and 0.337 for GABBI markers.

**Figure 3:**
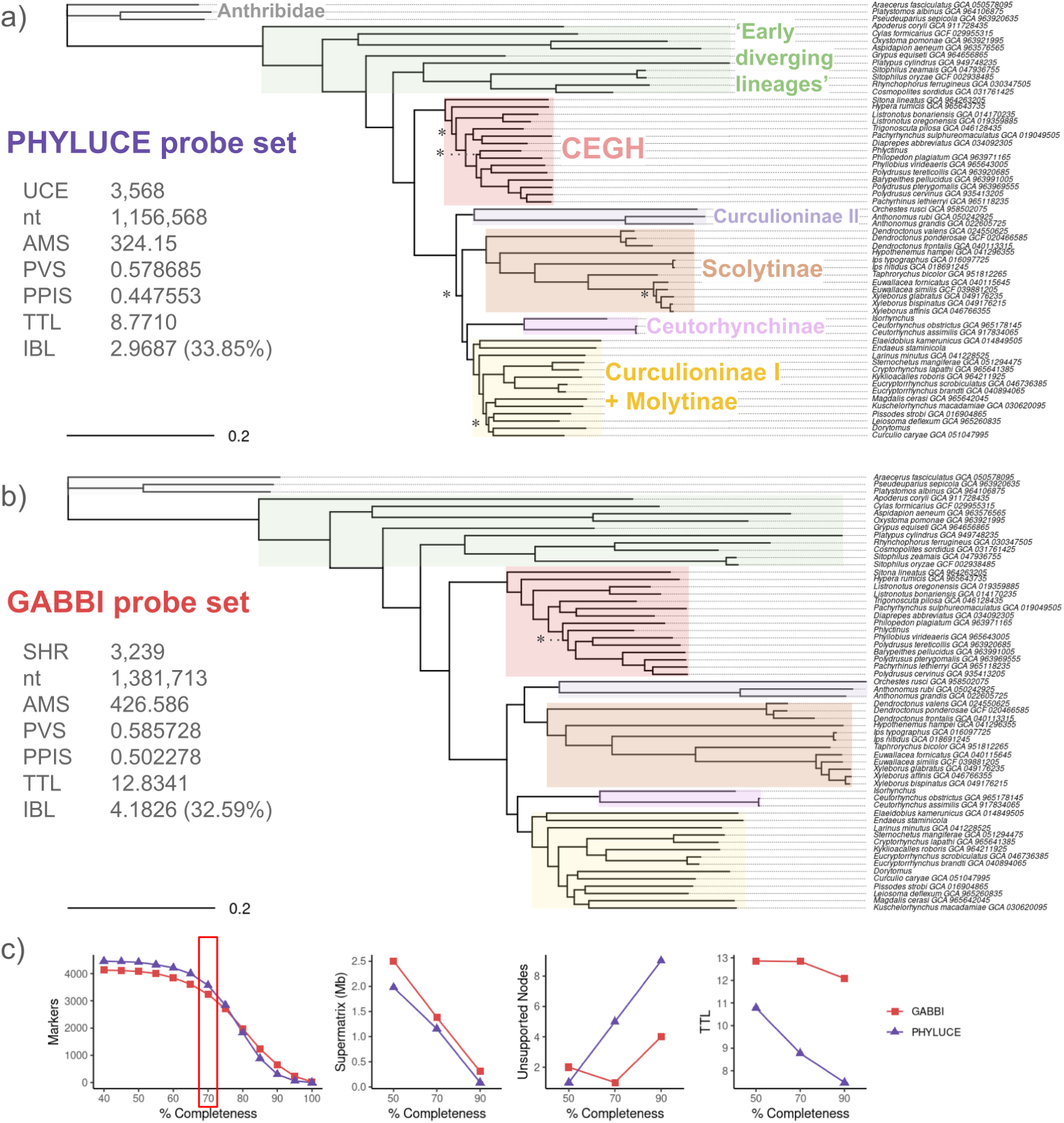
Scaled comparison between Curculionoidea phylogenomic trees obtained with PHYLUCE and GABBI probe sets. Both tree inferences were generated in a ML framework in IQTREE3 using the *-m MFP+MERGE* option and restricted to markers represented by at least 70% of total taxa. The scale is exactly the same between both trees to highlight differences of tree size. Unsupported nodes are marked with (*) if only UFBS fail or (**) if SH-aLRT and UFBS fall below 80 and 95%, respectively. Main weevil groups are shaded with different colors to better compare topologies and tree lengths. a) Species tree of Curculionoidea obtained following *in silico* target capture of PHYLUCE-derived markers. UCE = ultra-conserved elements, nt = nucleotides in the supermatrix, MMS = mean marker size (combining core and flanking regions), PVS = proportion of variable sites, PPIS = proportion of parsimony informative sites, TTL = total tree length, IBL = internal branch length and proportion to TTL. b) Species tree of Curculionoidea obtained following *in silico* target capture of GABBI-derived markers. SHR = shared homologous regions. c) Comparative statistics between PHYLUCE and GABBI-derived trees depending on supermatrix completeness (i.e. the minimum percentage of taxa required to keep a locus). The red rectangle indicates the supermatrix completeness represented.

#### Pachyrhynchini dataset

Only markers represented by more than 50% and 70% of all taxa in the dataset were kept for the concatenated phylogenomic analysis, as no dataset allowed the conservation of more than 90% of all taxa in any alignment. Supermatrix sizes range from 266,716 nt to 1,031,203 nt in the PHYLUCE dataset, and from 261,147 nt to 901,787 nt in the GABBI dataset. As in the *Curculionoidea* dataset, PHYLUCE-derived trees have more unsupported nodes and shorter branch lengths as GABBI-derived trees although they have comparable supermatrix sizes (Fig. S3-4; Table S3). At 70% completeness, PHYLUCE and GABBI-derived supermatrices were respectively 266,716 nt and 261,147 nt long and produced phylogenomic trees with 23 and 15 unsupported nodes, respectively, while total tree length was estimated to be 0.650 units long in the PHYLUCE-derived tree and 1.515 units long in the GABBI-derived tree (Fig. S4). In comparison, Van Dam et al. ’s custom UCE probe set provided 6,659 markers at 70% supermatrix completeness, resulting in a 1,181,120 nt long supermatrix but a total tree length of only 1.412 units long and 13 unsupported nodes (Fig. S5). Additionally, gene trees were computed with the GTR+G8 model for each marker allowed us to get a median value for the gamma function’s alpha parameter of 0.408 for PHYLUCE markers, 0.429 for GABBI markers and 0.689 for Van Dam et al.’s markers (Table S3).

### Available Coleoptera probe sets performances

Currently available Coleoptera UCE and Coleoptera AHE probe sets were tested at the same taxonomic scales and in the same methodological framework as above, except that the annotation of coding partitions could not be estimated for these probe sets. Resulting statistics are compared to those of GABBI-derived datasets that were generated again without codon annotation (Table 1). Overall, the Coleoptera UCE probe set is slightly outperformed by the Coleoptera AHE and GABBI probe sets at the taxonomic scales tested here. Despite a much smaller number of targeted loci, the Coleoptera AHE probe set has comparable values of average PVS, average PPIS, median alpha and branch lengths as the GABBI probe set. However, the number of unsupported nodes is higher with the Coleoptera AHE probe set and increases with supermatrix completeness, while staying at its lowest value with the GABBI probe set regardless of the supermatrix completeness (2 out of 57 bootstraps in the *Curculionoidea* dataset and 11-12 out of 78 bootstraps in the *Pachyrhynchini* dataset). In the *Curculionoidea* dataset, topologies inferred are very close between probe sets, but differ on the position of Ceuthorhynchinae, either as a sister group of the Curculioninae II*+*Scolytinae clade in all Coleoptera AHE trees and the 50% completeness Coleoptera UCE tree, or as a sister group of the Curculioninae I+Molytinae clade in all GABBI-derived trees and the 70% completeness Coleoptera UCE tree (Fig. S6-8; Supplementary Material on Zenodo).

Interestingly, *in silico* target capture using the GABBI probe set on already generated AHE data can be achieved following the same workflow used for capturing GABBI markers on Van Dam et al’s dataset, using raw read data from Haran et al. (2023). On 30 individuals tested, an average number of 558 markers (out of 4,255) have been successfully recovered from raw reads, ranging from 229 to 1,336 markers per individual. The 50% completeness supermatrix obtained following alignments and cleaning is 1,197,403 nt long and successfully positioned all 30 taxa in the resulting phylogenomic tree, with on average 90.91% gap/ambiguity (Fig. S9).

**Table 1:** Phylogenomic statistics following *in silico* target capture of currently available Coleoptera probe sets and the new GABBI weevil probe set. The highest values of a given taxonomic dataset are represented in bold.

| Dataset | Probe set | Total number of clean markers | Average marker size (bp.) | Average PVS | Number of PIS | Average PPIS |
| --- | --- | --- | --- | --- | --- | --- |
| *Curculionoidea* Superfamily | Weevil GABBI | **4,136** | **697.31** | **0.659** | **1,543,634** | **0.546** |
|  | Coleoptera UCE | 1065 | 507.595 | 0.564 | 232,272 | 0.433 |
|  | Coleoptera AHE | 747 | 656.068 | 0.633 | 254,784 | 0.521 |
| *Pachyrhynchini* Tribe | Weevil GABBI | **3,759** | **357.037** | 0.401 | **316,036** | 0.240 |
|  | Coleoptera UCE | 916 | 308.678 | 0.292 | 42,764 | 0.155 |
|  | Coleoptera AHE | 590 | 349.551 | **0.416** | 51,627 | **0.252** |

| Median<br>alpha<br>value | 50%<br>completeness<br>Supermatrix<br>size (nt.) | Number of<br>unsupported<br>nodes (50%) | Total tree<br>length<br>(50%) | Internal<br>branch length<br>(50%) | 70%<br>completeness<br>Supermatrix<br>size (nt.) |
| --- | --- | --- | --- | --- | --- |
| **0.33575** | **2,643,888** | **2** | 12.8084 | **3.9918** | **1,620,294** |
| 0.303 | 229,616 | 7 | 10.4238 | 3.2797 | 79,286 |
| 0.317 | 367,108 | 3 | **13.4507** | 3.9307 | 191,756 |
| 0.4374 | **925,886** | **11** | 1.8366 | 0.4838 | **295,093** |
| 0.38185 | 114,699 | 30 | 1.0759 | 0.2786 | 12,064 |
| **0.4444** | 145,158 | 20 | **1.9449** | **0.4975** | 48,865 |

| Number of<br>unsupported<br>nodes (70%) | Total<br>tree<br>length<br>(70%) | Internal<br>branch<br>length<br>(70%) | 90%<br>completeness<br>Supermatrix<br>size (nt.) | Number of<br>unsupported<br>nodes<br>(90%) | Total tree<br>length<br>(90%) | Internal<br>branch<br>length<br>(90%) |
| --- | --- | --- | --- | --- | --- | --- |
| **2** | 12.5851 | **4.0707** | **413,667** | **2** | 11.5832 | 3.894 |
| 11 | 8.6634 | 2.8242 | 1,975 | 36 | 4.4924 | 1.5913 |
| 6 | **13.3858** | 4.0454 | 58,250 | 11 | **16.8453** | **5.2952** |
| **12** | **1.597** | 0.4216 | 0 | - | - | - |
| 53 | 0.649 | 0.1712 | 0 | - | - | - |
| 32 | 1.5516 | **0.4285** | 0 | - | - | - |

PVS = Proportion of variable sites, PIS = Parsimony informative sites, PPIS = Proportion of PIS.

### In vitro *target capture*

The GABBI-derived probe set was synthesized and tested in vitro on 12 weevil samples. Sample metadata, DNA extraction statistics and phylogenomic statistics using this probe set are summarised in Table 2. An average of 6,355,037 paired-end reads (146 bp long) were obtained following read cleaning. The iterative assemblies recovered an average of 3,628 SHR (85% of total markers), ranging from 3,168 to 3,825 SHR (74% to 90%). Their length before aligning them with the *Curculionoidea dataset* were on average 695 to 1,189 bp long. Following alignment cleaning, core and flanking regions were on average 186.955 nt and 243.054 nt long, respectively, adding up to a 70% completeness supermatrix of 2,159,524 nt with the *Curculionoidea dataset* and an average amount of missing data of 25.95% (Table 2). Their phylogenomic position within Curculionoidea is shown in figure 4.

**Table 2:**
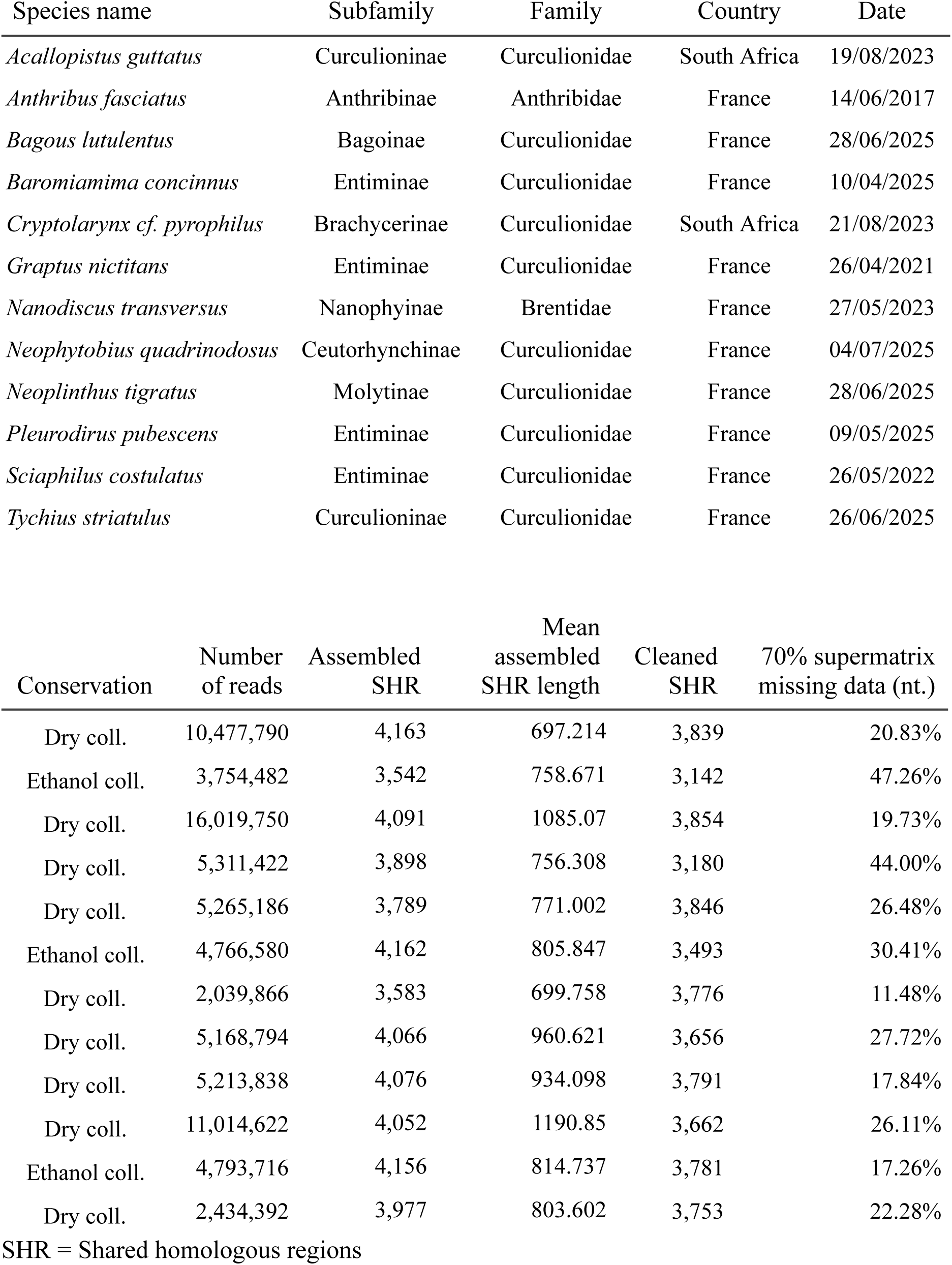
DNA extraction and phylogenomic statistics of samples processed with the GABBI-derived weevil probe set.

**Figure 4:**
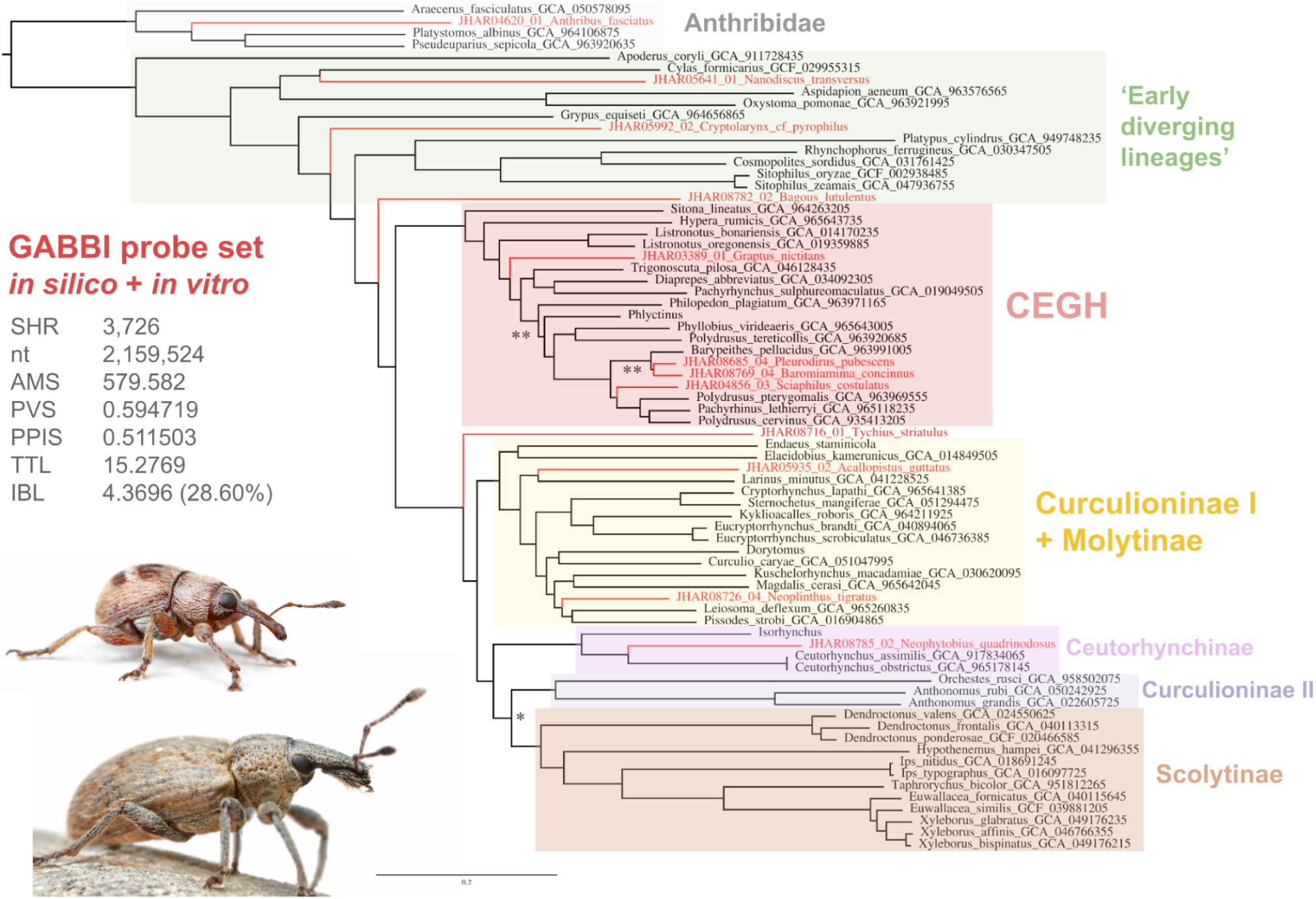
Phylogenomic inference using GABBI weevil probe set combining the *Curculionoidea dataset* with *in vitro* target capture specimens. The inferred tree results from a ML framework using IQTREE3 on markers represented by at least 70% of total taxa. Evolution models were computed with the *-m MFP+MERGE* option. Unsupported nodes are marked with (*) if only UFBS fail or (**) if SH-aLRT and UFBS fall below 80 and 95%, respectively. Main weevil groups are shaded as in figure 3. Specimens obtained from *in silico* target capture of the *Curculionoidea dataset* are shown on dark branches whereas specimens obtained from *in vitro* target capture are shown on red branches. SHR = shared homologous regions, nt = nucleotides in the supermatrix, MMS = mean marker size (combining core and flanking regions), PVS = proportion of variable sites, PPIS = proportion of parsimony informative sites, TTL = total tree length, IBL = internal branch length and proportion to TTL. Credit for pictures: top: *Nanodiscus transversus*, Martin Galli; bottom: *Graptus nictitans*, Benjamin Zelvelder.

## DISCUSSION

### Comparison of GABBI and PHYLUCE Probe Design Approaches

Designing probe sets for target capture strategies has become an essential part in modern phylogenomics of non-model species. Although a great variation in marker selection methods have been proposed by the scientific community (e.g. Lemmon et al. 2012; Faircloth 2017; Hutter et al. 2022; Ball et al. 2023), none have yet taken advantage of recent genome-alignment methods to identify shared genomic markers. Using weevils as models (Coleoptera, Curculionoidea), we tested GABBI performance to design probe sets against the widely used and open-source PHYLUCE pipeline (Faircloth 2016). Differences between both methods can already be drawn during probe design, as PHYLUCE provided fewer markers in the final set of targeted loci than GABBI when imposing a minimum threshold of 90% shared taxa and had to be reduced to a threshold of 80%. Both methods provided around 320k extent and ancestral sequences of targeted loci, but the GABBI probe set ended up with twice as much probe sequences as the PHYLUCE probe set after applying the exact same clustering method (Fig. 1). This means that there were not as many redundant probes in the GABBI probe set as in the PHYLUCE one, suggesting that GABBI-dervied markers are more variable and need more probe sequences to map their variability.

Nonetheless, this difference in variability did not affect the number of loci recovered from *in silico* nor *in vitro* target capture (Fig. 4; Table S2), but it probably explains why we manage to get GABBI-derived markers with, on average, significantly more parsimony informative sites within targeted (core) regions than PHYLUCE-derived ones (Fig. 2). This results in phylogenomic trees with longer branches and stronger support in GABBI-derived trees, which can thus be considered as more phylogenetically resolutive, especially at higher supermatrix completeness values (expressed as a minimum percentage of taxa required to keep a locus; Fig. 3c, S1-2). Importantly, this tendency is not challenged by a shallower taxonomic scale, as it shows an even more contrasted difference at the tribe scale in favor of GABBI-derived markers (Fig. S3-4). However, although we expected the alpha parameter of the gamma distribution to be a good predictor of phylogenetic signal (Yang 1996; Allio et al. 2024), our results did not show notable differences in the distributions of any set of markers (Table S3). To avoid any bias linked to marker selection and keep as much data as possible, we chose not to rely on this variable for marker selection or discussion, but that would be a possibility if a much bigger dataset needed to be cropped. Interestingly, markers selected by PHYLUCE seem to match with the definition of “ultraconserved elements” as sequences that are not necessarily coding but are surrounded by more variable flanking data (Faircloth et al. 2012). However, GABBI-derived markers cannot truly be considered as ultraconserved elements and should rather be identified as the newly proposed term of shared homologous regions (SHR) for their greater level of inner variability. The great proportion of coding regions among GABBI markers might, however, only be a consequence of better genomic alignments within coding regions but should not be intrinsic to SHR.

### Performance of the new GABBI-derived Weevil Probe Set

Compared to currently available target capture kits that could be used for weevil phylogenomics, namely the Coleoptera UCE (Faircloth 2017) or Coleoptera AHE (Haddad et al. 2018), our newly designed set of weevil probes provides an essential alternative that outperforms them in several aspects (Table 1). Although AHE markers were selected at the Coleoptera scale, they often show similar statistics as GABBI markers (Table 1). However, their limited number and low percentage of recovery when used at the weevil scale (about ⅓ of all 941 AHE were represented in Shin et al. 2018; Haran et al. 2023b; or Li et al. 2023) emphasize the benefit of the new GABBI-derived set of markers for weevils that can recover more than 10 times more markers per individual without losing marker quality.

Although the aim of this study is not to formally address a stable phylogeny of weevils, the phylogenomic inference of the Curculionoidea tree with both *in silico* and *in vitro* samples appears very robust and agrees with the most recent molecular analyses in weevils (Fig. 4; Haran et al. 2023b; Li et al. 2023). Most importantly, the weevil specific probe kit newly designed in this study provided excellent levels of marker recovery for *in vitro* samples, whether they were dry-mounted or conserved in alcohol (Table 2). Future work will be needed to assess the efficiency of this probe set for older museum samples, but it should perform better than usual UCE kits by providing more numerous and resolutive targeted core regions, as degraded DNA of museum samples often results in shorter contigs with reduced or absent flanking regions, potentially approaching core-only sequences (McCormack et al. 2016; Smith et al. 2020). GABBI-derived probe sets should thus provide a new set of alternative markers for taxonomic groups facing similar issues with probe sets designed at large taxonomic scales.

Some authors already circumvented this issues by designing specific probe sets for the taxa they studied (e.g. Gustafson et al. 2023; Van Dam et al. 2023; outside of Coleoptera: Ufimov et al. 2021 in plants, Chen et al. 2025 in mollusca). By greatly restricting the taxonomic scale at which the probe design is conducted, authors are able to get much more numerous loci and excellent levels of phylogenetically informative sites. However, such kits are often used for the sake of one or few studies, making them much more expensive and less useful for the scientific community.

Interestingly, the very large custom set of tribe-specific UCE designed by Van Dam et al. (2023) for Pachyrhynchini seems to hold much more phylogenetic information than other probe sets tested in this study given its greater number of markers, parsimony informative sites and bigger alpha value, but the resulting species tree is not longer than GABBI-derived and AHE-derived trees and has a comparable number of unsupported nodes as the GABBI-derived tree (Table S3). Thus, the GABBI approach seems to reach close performances to these custom probe designs while targeting a more diverse set of taxa.

### Benefits of a Genome Alignment based Bait Inference

The wide taxonomic range at which GABBI-derived markers perform well phylogenetically have several explanations. First, genome alignments involve a less stringent marker selection as read alignments. Although identifying shared genomic regions with genome alignments was expected to be more conservative and provide less candidate loci (Faircloth 2017), recent methods like Progressive Cactus proved to provide as many (if not more) conserved loci and hold a greater level of variation that isn’t allowed by a stringent mapping strategy. Second, as genome alignments don’t rely on a base taxon or reference genome, we avoid multiple biases linked to base genome selection that could be caused by genome quality, genome size, large individual structural variants or simply the highly specific gene-pool of a given species (as pointed out by Gustafson et al. 2019). The genomic coverage between one taxon and the reference is also expected to decrease rapidly with taxonomic distance (Armstrong et al. 2020). By avoiding this ‘single-reference’ bias, the GABBI pipeline provides a more diverse and universal set of markers (i.e. shared by a greater number of taxa) that showed to be more robust to an increasing stringency on missing data (Fig. 3; S4). Third, instead of inevitably reducing the number of shared markers when adding an increasing amount of distantly related taxa in the probe design, because some of them become duplicated in some lineages or come missing in too many, allowing a maximum of one duplication event during the early stages of the probe design greatly increases the number of candidate loci that will be tested at a larger scale. Even though this implies that some markers can be duplicated in a given lineage (and must be removed before phylogenomic analysis, as we have done with marker blacklisting), the same markers may not be duplicated in another lineage and used safely in the latter, providing more numerous targeted loci at a larger taxonomic scale, making the probe set adaptable to many subsamples of the initially targeted taxonomic range. The probability of selecting ‘false negative’ single-copy markers (paralogous loci) should decrease rapidly with an increasing number of genomes in the genomic alignment and a large number of additional genomes to test the temporary probe set on.

### Conclusion

In all taxonomic scales and supermatrix completeness tested, GABBI-derived markers designed for weevils provide better phylogenetic resolution in targeted regions than what we would obtain with a mapping-based method of probe design. Using this method, we provide the first weevil-specific probe set for target capture that enhances the marker recovery and phylogenomic resolution expected with currently available probe sets designed on larger taxonomic scales. Thanks to big consortiums that fuel high quality genome sequencing efforts, it is and will become increasingly easier to get high quality genome alignments at the basis of this probe design method. While target capture methods are becoming essential for their usefulness in museomics and ultradiverse non-model species, the GABBI pipeline provides a way of enhancing probe design by combining it with advances in genome sequencing. By making the GABBI approach easily accessible to anyone as an open-source, turnkey tool available on github (https://github.com/bjzelvelder/GABBI.git), we hope we can make targeted capture research move forward alongside genomic advances, as it remains the most effective way to apprehend the evolution of highly diverse, non-model taxa.

## FUNDING ACKNOWLEDGMENT

This work was supported by the French National Research Agency (grant number ANR-24-CE02-0309); and by the Santé des Plantes et Environnement departement of INRAe (project name: WeBait).

## ACKNOWLEDGMENTS

We would like to thank Jacob Enk for helping us design and optimize the probe sets; Frédéric Delsuc and Emmanuel Douzery for their insightful discussions during the development of the GABBI pipeline; and Nicolas Zelvelder for his help in designing the GABBI logo. We acknowledge the genotoul bioinformatics platform Toulouse Occitanie (Bioinfo Genotoul, https://doi.org/10.15454/1.5572369328961167E12) for providing computing and storage resources. The following services are acknowledged for access and collecting permits: the Western Cape Nature Conservation Board (Permit No. CN44-30-4229; Republic of South Africa), the Cape Research Centre, South African National Parks (Permit No. CRC/2019-2020/012–2012/V1) and the Mercantour National Park (France). This is the contribution ISEM #xxx of the Institut des Sciences de l’Evolution de Montpellier.

## DATA AVAILABILITY STATEMENT

Once submitted, newly generated genomic assemblies and raw target capture data will be available on Dryad at xxx. Scripts and Supplementary Material will be available on Zenodo at xxx.

