## Supplementary material for "A new method based on genome alignments provides a highly resolutive target enrichment set for weevils (Coleoptera, Curculionoidea)": Fig. S

DOI:

**Figure S1:** Scaled comparison between Curculionoidea phylogenomic trees obtained with PHYLUCE and GABBI probe sets. (50% Supermatrix completeness)

**Figure S2:** Scaled comparison between Curculionoidea phylogenomic trees obtained with PHYLUCE and GABBI probe sets. (90% Supermatrix completeness)

**Figure S3:** Scaled comparison between Pachyrhynchini phylogenomic trees obtained with PHYLUCE and GABBI probe sets. (50% Supermatrix completeness)

**Figure S4:** Scaled comparison between Pachyrhynchini phylogenomic trees obtained with PHYLUCE and GABBI probe sets. (70% Supermatrix completeness)

**Figure S5:** Phylogenomic tree obtained with Van Dam et al. 's (2023) probe set on the Pachyrhynchini dataset.

**Figure S6:** Phylogenomic tree obtained with the GABBI probe set on the Curculionoidea dataset for comparison with existing Coleoptera target capture sets.

**Figure S7:** Phylogenomic tree obtained with the Coleoptera UCE probe set on the Curculionoidea dataset.

**Figure S8:** Phylogenomic tree obtained with the Coleoptera AHE probe set on the Curculionoidea dataset.

**Figure S9:** Phylogenomic tree obtained with the GABBI probe set on the Curculionoidea dataset and Haran et al. 's (2023) dataset obtained from Coleoptera AHE probe set.

Dryad repository:

Zenodo repository:

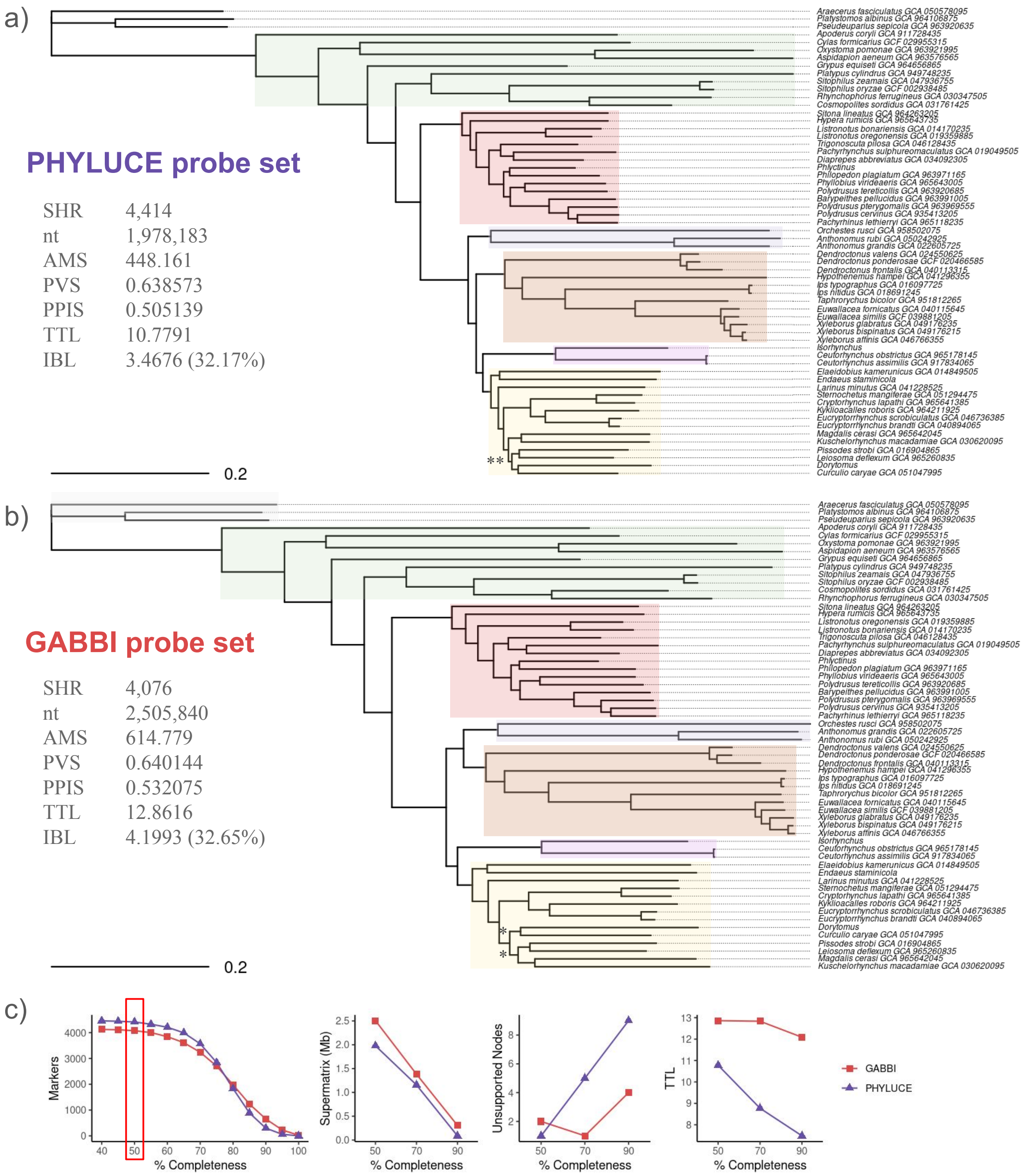


**Figure S1: Scaled comparison between Curculionoidea phylogenomic trees obtained with PHYLUCE and GABBI probe sets.** Both tree inferences were generated in a maximum likelihood framework in IQTREE3 using the *-m MFP+MERGE* option and restricted to markers represented by at least 50% of total taxa. The scale is exactly the same between both trees to highlight differences of tree size. Unsupported nodes are marked with (*) if only UFBS fail or (**) if SH-aLRT and UFBS fall below 80 and 95%, respectively. Main weevil groups are shaded with different colors to better compare topologies and tree lengths. a) Species tree of Curculionoidea obtained following in silico target capture of PHYLUCE-derived markers. UCE = ultra-conserved elements, nt = nucleotides in the supermatrix, MMS = mean marker size (combining core and flanking regions), PVS = proportion of variable sites, PPIS = proportion of parsimony informative sites, TTL = total tree length, IBL = internal branch length and proportion to TTL. b) Species tree of Curculionoidea obtained following *in silico* target capture of GABBI-derived markers. SHR = shared homologous regions. c) Comparative statistics between PHYLUCE and GABBI-derived trees depending on supermatrix completeness (i.e. the minimum percentage of taxa required to keep a locus). The red rectangle indicates the supermatrix completeness represented.

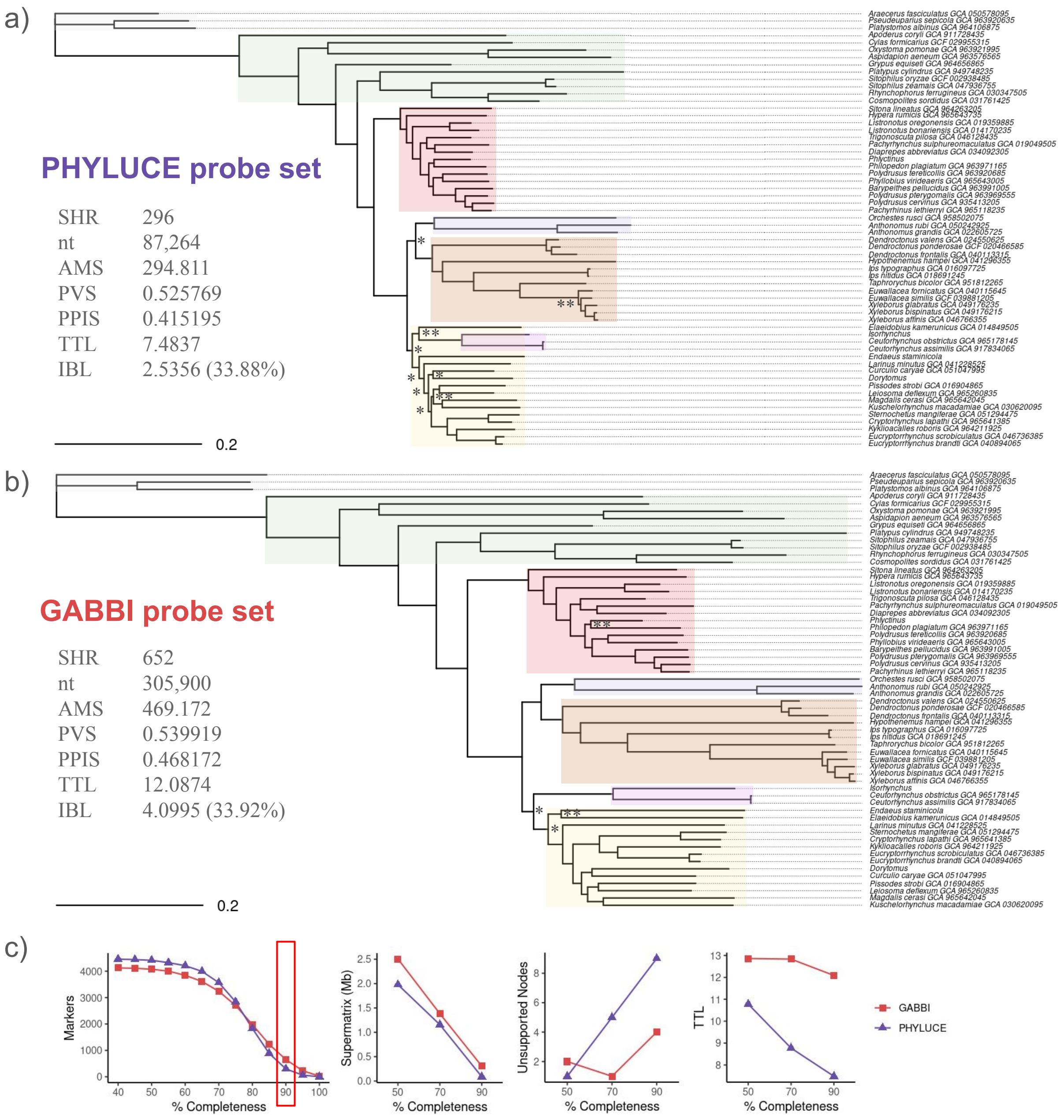


**Figure S2: Scaled comparison between Curculionoidea phylogenomic trees obtained with PHYLUCE and GABBI probe sets.** Both tree inferences were generated in a maximum likelihood framework in IQTREE3 using the *-m MFP+MERGE* option and restricted to markers represented by at least 90% of total taxa. The scale is exactly the same between both trees to highlight differences of tree size. Unsupported nodes are marked with (*) if only UFBS fail or (**) if SH-aLRT and UFBS fall below 80 and 95%, respectively. Main weevil clades are shaded with different colors to better compare topologies and tree lengths. a) Species tree of Curculionoidea obtained following in silico target capture of PHYLUCE-derived markers. UCE = ultra-conserved elements, nt = nucleotides in the supermatrix, MMS = mean marker size (combining core and flanking regions), PVS = proportion of variable sites, PPIS = proportion of parsimony informative sites, TTL = total tree length, IBL = internal branch length and proportion to TTL. b) Species tree of Curculionoidea obtained following *in silico* target capture of GABBI-derived markers. SHR = shared homologous regions. c) Comparative statistics between PHYLUCE and GABBI-derived trees depending on supermatrix completeness (i.e. the minimum percentage of taxa required to keep a locus). The red rectangle indicates the supermatrix completeness represented.

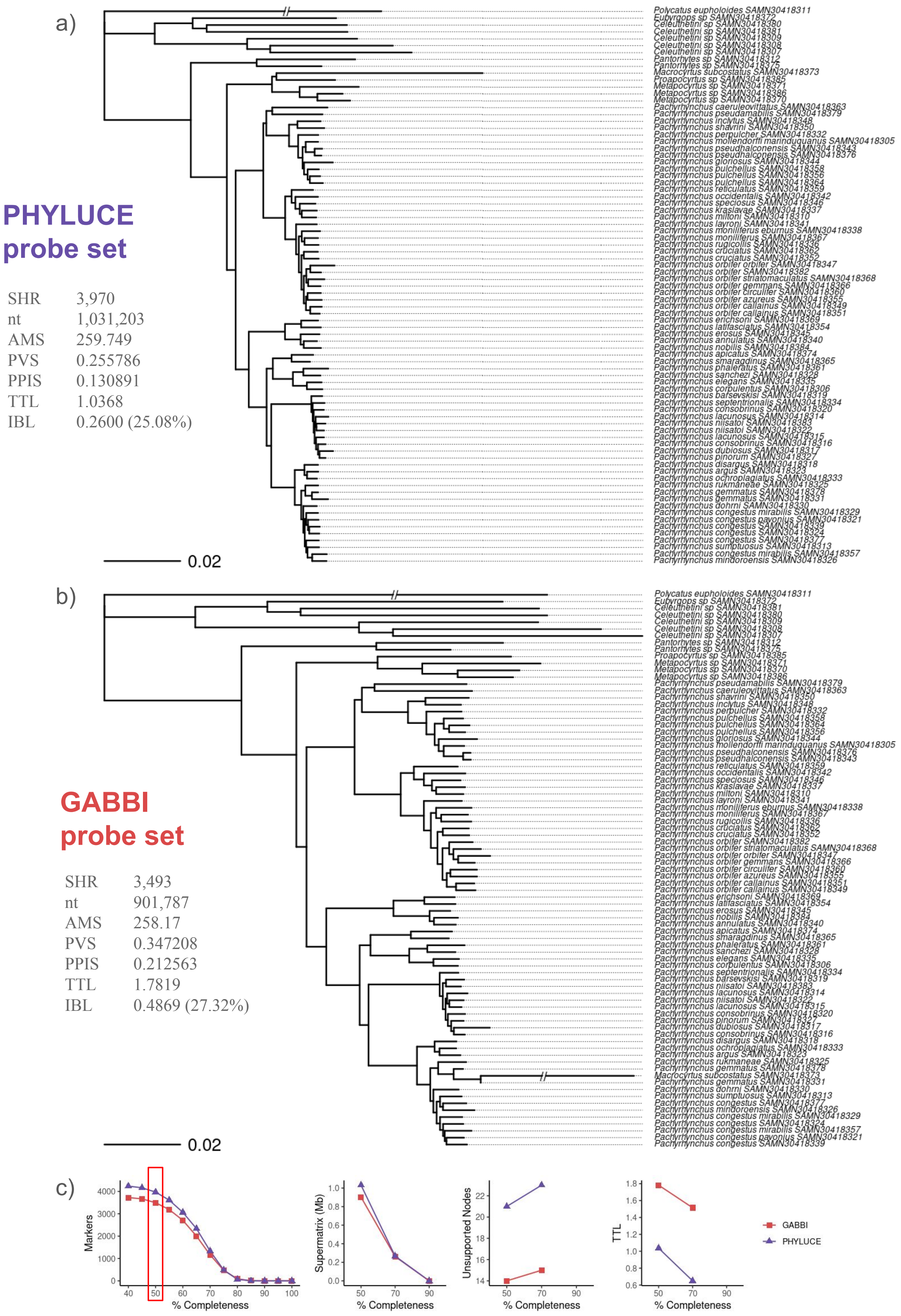


**Figure S3: Scaled comparison between Pachyrhynchini phylogenomic trees obtained with PHYLUCE and GABBI probe sets.** Both tree inferences were generated in a maximum likelihood framework in IQTREE3 using the *-m MFP+MERGE* option and restricted to markers represented by at least 50% of total taxa. The scale is exactly the same between both trees to highlight differences of tree size. a) Species tree of Pachyrhynchini obtained following in silico target capture of PHYLUCE-derived markers. UCE = ultra-conserved elements, nt = nucleotides in the supermatrix, MMS = mean marker size (combining core and flanking regions), PVS = proportion of variable sites, PPIS = proportion of parsimony informative sites, TTL = total tree length, IBL = internal branch length and proportion to TTL. b) Species tree of Pachyrhynchini obtained following *in silico* target capture of GABBI-derived markers. SHR = shared homologous regions. c) Comparative statistics between PHYLUCE and GABBI-derived trees depending on supermatrix completeness (i.e. the minimum percentage of taxa required to keep a locus). The red rectangle indicates the supermatrix completeness represented.

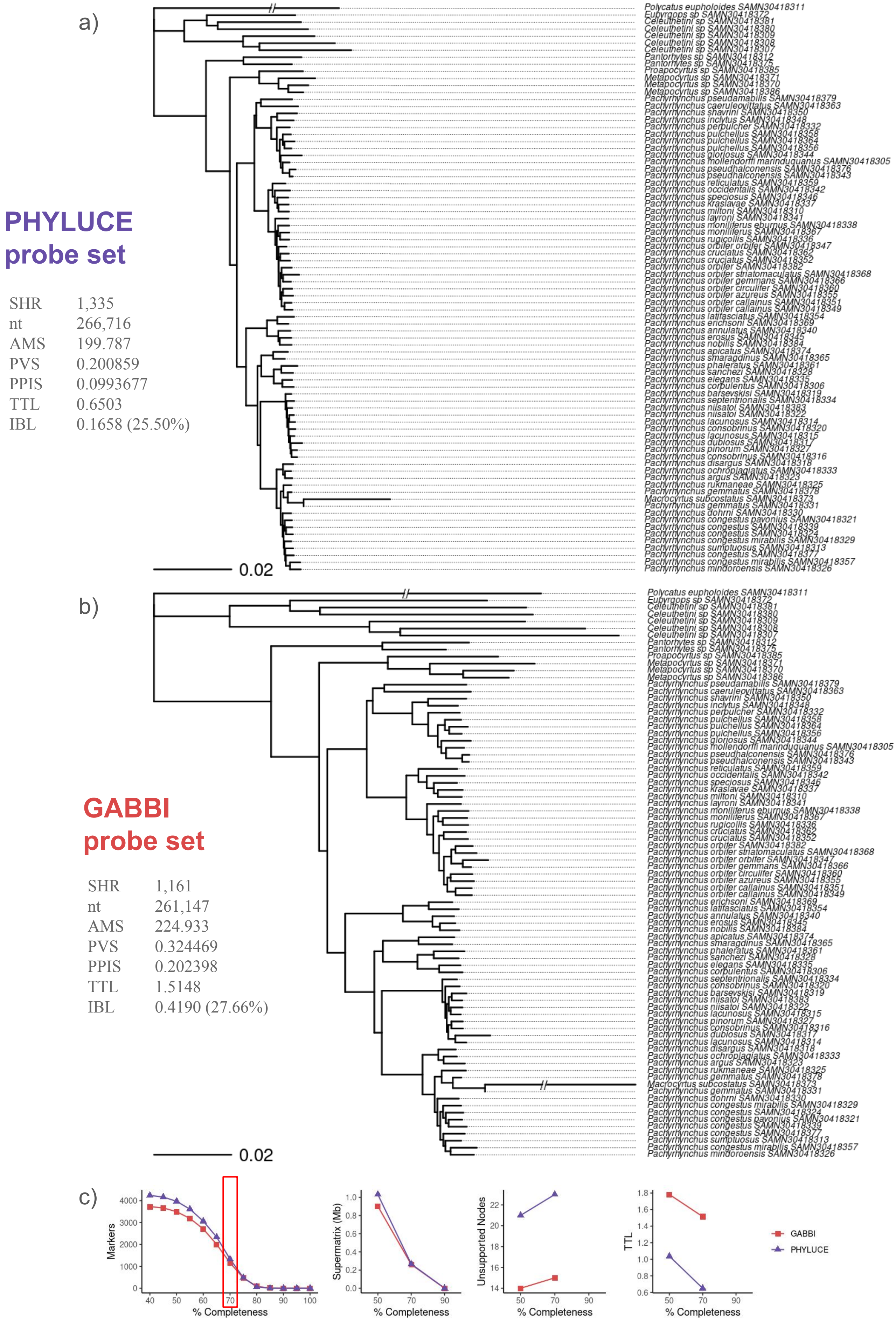


**Figure S4: Scaled comparison between Pachyrhynchini phylogenomic trees obtained with PHYLUCE and GABBI probe sets.** Both tree inferences were generated in a maximum likelihood framework in IQTREE3 using the *-m MFP+MERGE* option and restricted to markers represented by at least 70% of total taxa. The scale is exactly the same between both trees to highlight differences of tree size. a) Species tree of Pachyrhynchini obtained following in silico target capture of PHYLUCE-derived markers. UCE = ultra-conserved elements, nt = nucleotides in the supermatrix, MMS = mean marker size (combining core and flanking regions), PVS = proportion of variable sites, PPIS = proportion of parsimony informative sites, TTL = total tree length, IBL = internal branch length and proportion to TTL. b) Species tree of Pachyrhynchini obtained following *in silico* target capture of GABBI-derived markers. SHR = shared homologous regions. c) Comparative statistics between PHYLUCE and GABBI-derived trees depending on supermatrix completeness (i.e. the minimum percentage of taxa required to keep a locus). The red rectangle indicates the supermatrix completeness represented.

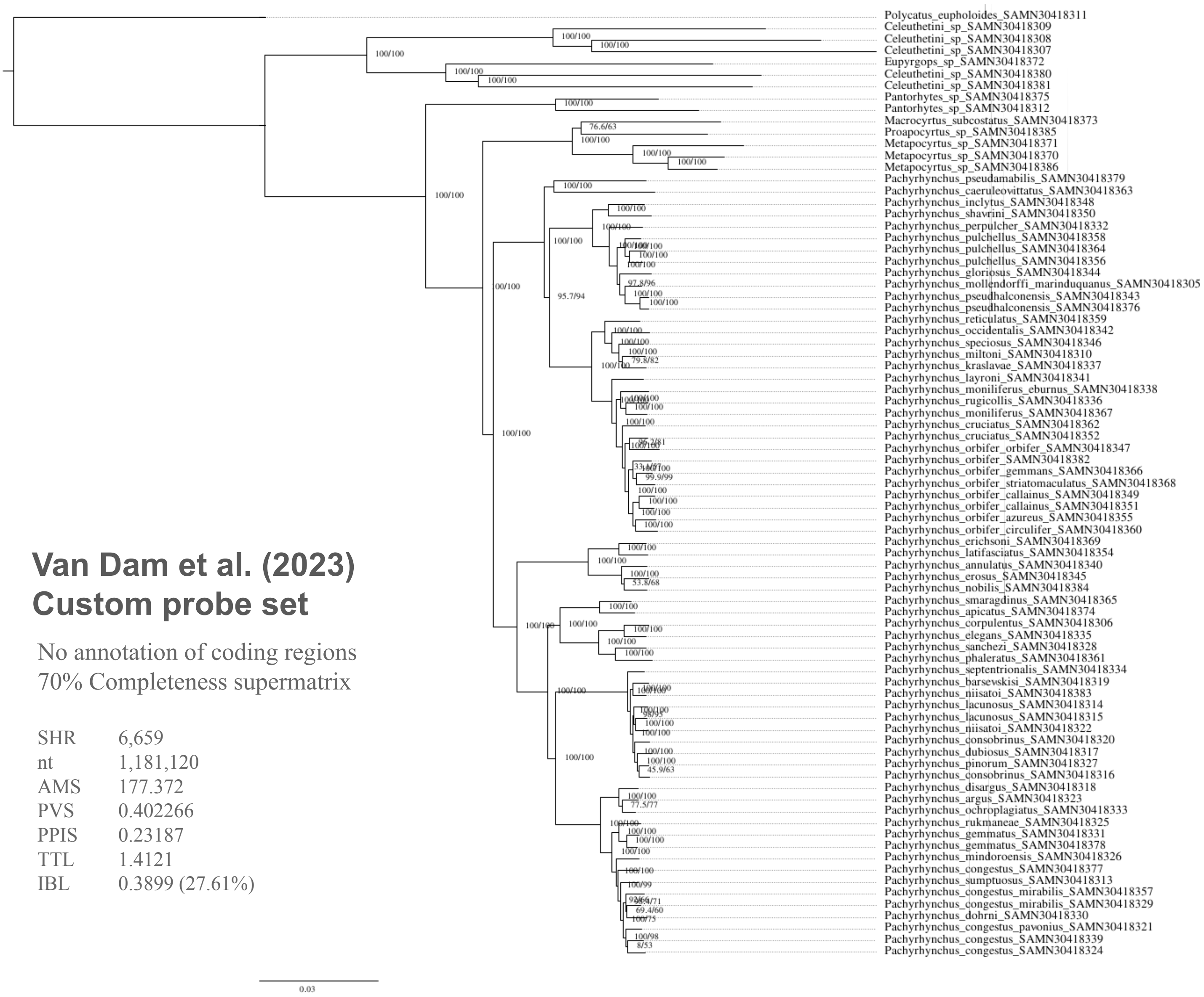


**Figure S5: Phylogenomic tree obtained with Van Dam et al. 's (2023) probe set on the *Pachyrhynchini dataset*.** The tree was inferred in a maximum likelihood framework in IQTREE3 using the *-m MFP+MERGE* option and restricted to markers represented by at least 70% of total taxa. Partitions did not take into account codon positions. UCE = ultra-conserved elements, nt = nucleotides in the supermatrix, MMS = mean marker size (combining core and flanking regions), PVS = proportion of variable sites, PPIS = proportion of parsimony informative sites, TTL = total tree length, IBL = internal branch length and proportion to TTL.

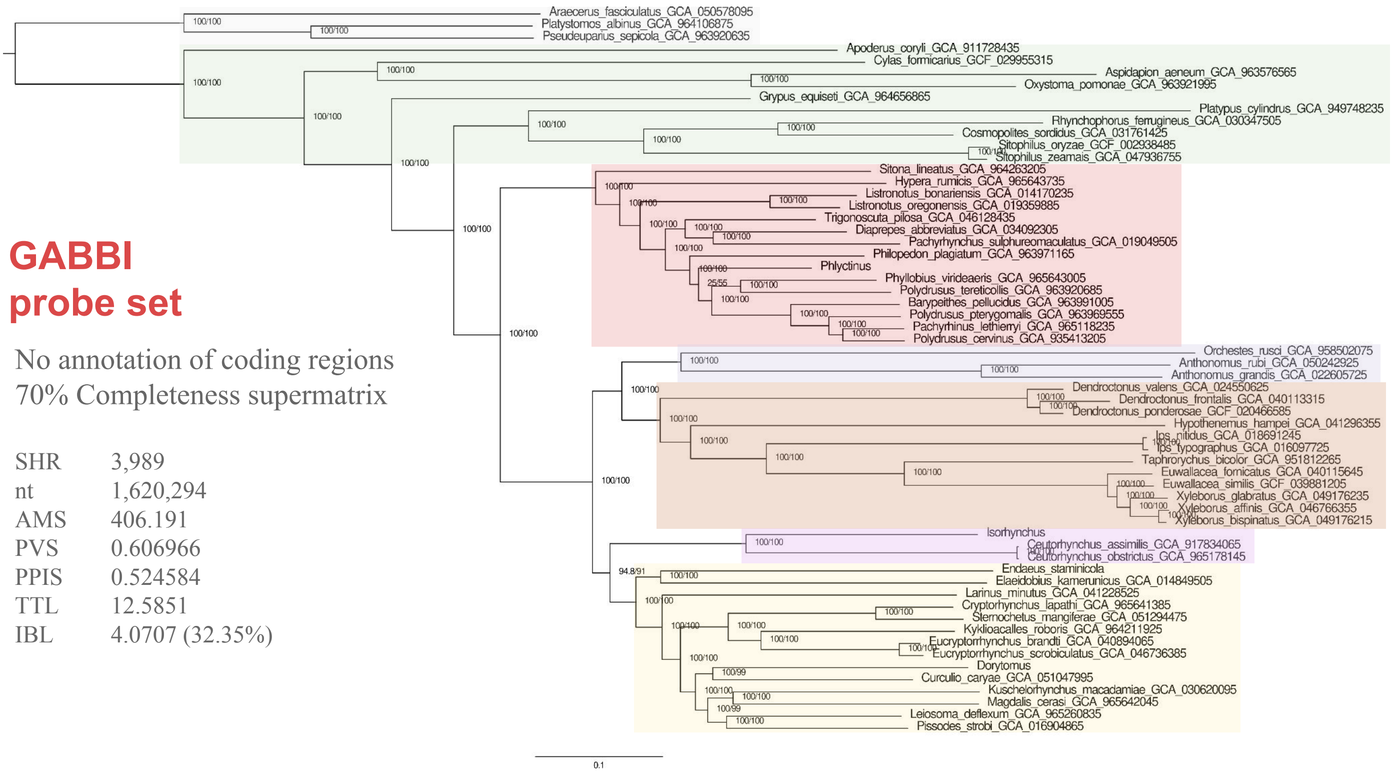


**Figure S6: Phylogenomic tree obtained with the GABBI probe set on the *Curculionoidea dataset* for comparison with existing Coleoptera target capture sets.** The tree was inferred in a maximum likelihood framework in IQTREE3 using the *-m MFP+MERGE* option and restricted to markers represented by at least 70% of total taxa. Partitions did not take into account codon positions. Main weevil groups are shaded as in figure 3. SHR = shared homologous regions, nt = nucleotides in the supermatrix, MMS = mean marker size (combining core and flanking regions), PVS = proportion of variable sites, PPIS = proportion of parsimony informative sites, TTL = total tree length, IBL = internal branch length and proportion to TTL.

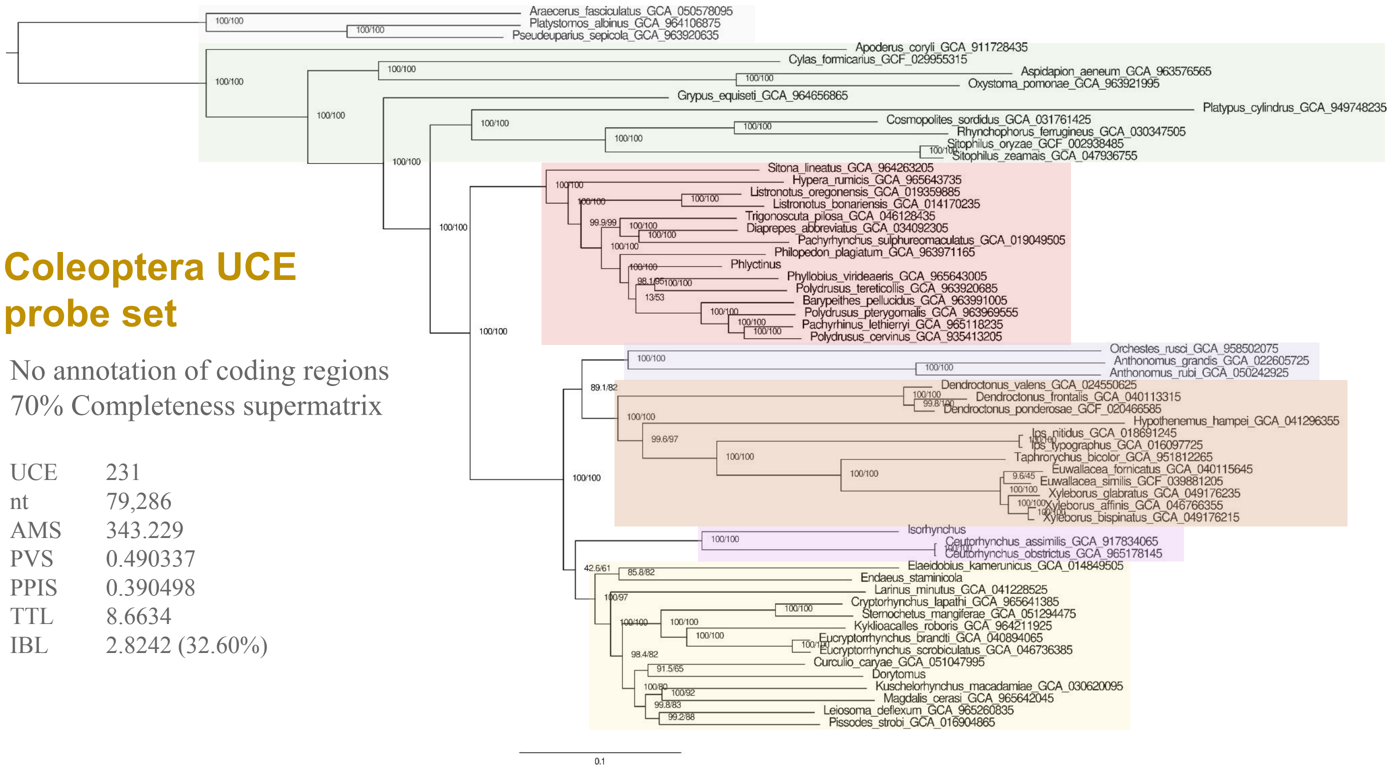


**Figure S7: Phylogenomic tree obtained with the Coleoptera UCE probe set on the *Curculionoidea dataset*.** The tree was inferred in a maximum likelihood framework in IQTREE3 using the *-m MFP+MERGE* option and restricted to markers represented by at least 70% of total taxa. Partitions did not take into account codon positions. Main weevil groups are shaded as in figure 3. SHR = shared homologous regions, nt = nucleotides in the supermatrix, MMS = mean marker size (combining core and flanking regions), PVS = proportion of variable sites, PPIS = proportion of parsimony informative sites, TTL = total tree length, IBL = internal branch length and proportion to TTL.

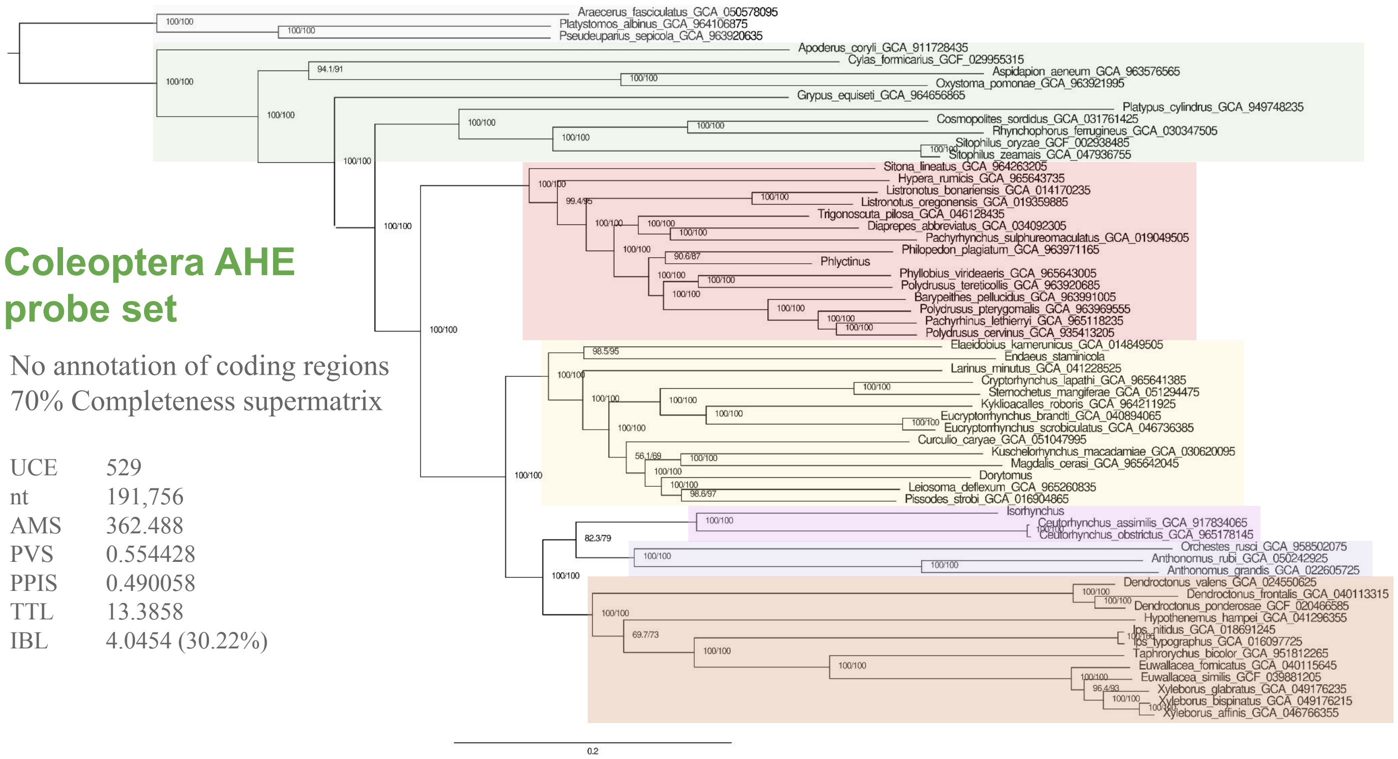


**Figure S8: Phylogenomic tree obtained with the Coleoptera AHE probe set on the *Curculionoidea dataset*.** The tree was inferred in a maximum likelihood framework in IQTREE3 using the *-m MFP+MERGE* option and restricted to markers represented by at least 70% of total taxa. Partitions did not take into account codon positions. Main weevil groups are shaded as in figure 3. SHR = shared homologous regions, nt = nucleotides in the supermatrix, MMS = mean marker size (combining core and flanking regions), PVS = proportion of variable sites, PPIS = proportion of parsimony informative sites, TTL = total tree length, IBL = internal branch length and proportion to TTL.

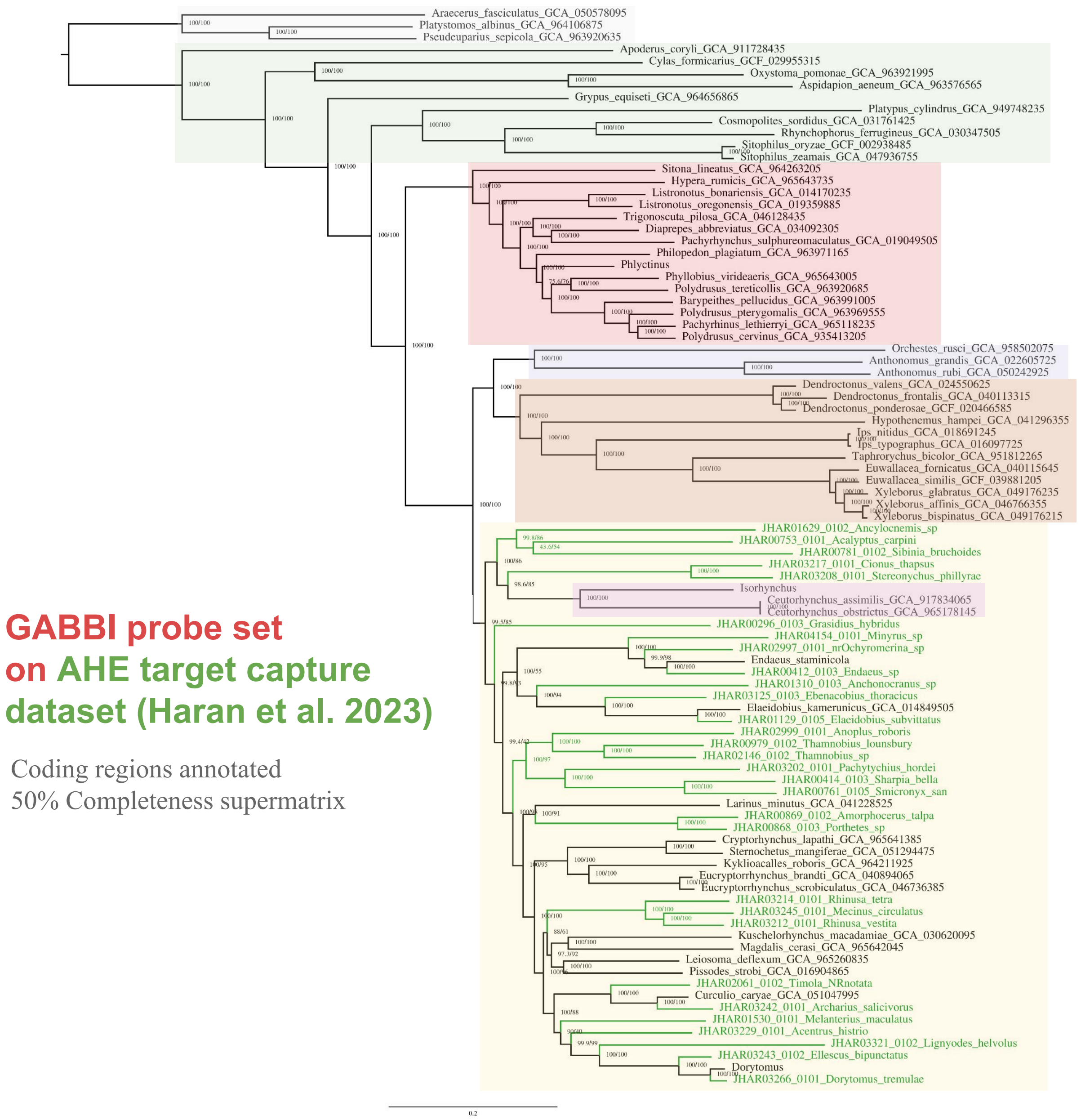


**Figure S9: Phylogenomic tree obtained with the GABBI probe set on the *Curculionoidea dataset* and Haran et al. 's (2023) dataset obtained from Coleoptera AHE probe set.** The tree was inferred in a maximum likelihood framework in IQTREE3 using the *-m MFP+MERGE* option and restricted to markers represented by at least 50% of total taxa. Main weevil groups are shaded as in figure 3.
